# Activation of IRF7/ISG15 axis in microglia inhibits NLRP3 Expression and Improves the Prognosis of Ischemia/Reperfusion in Mice

**DOI:** 10.1101/2025.11.08.687338

**Authors:** Hao Yan, Xiangyang Cao, Meng Ma, Qianxiong He, Feiyang Li, Ruxiang Xu, Yangyang Wang, Yi Zheng, Jing Yang, Ying Xiao

## Abstract

Ischemic stroke is a leading cause of disability and mortality, with neuroinflammation playing a key role in post-stroke injury. The molecular mechanisms remain incompletely defined. This study explored the functions of IRF7 and its downstream target ISG15 in stroke-associated neuroinflammation and prognosis of ischemic stroke. Bioinformatic analysis of transcriptomic datasets from microglia and ischemic brain tissues identified both molecules as hub genes. Their expression was validated in a mouse transient middle cerebral artery occlusion (tMCAO) model and in microglial cultures exposed to oxygen-glucose deprivation/reoxygenation (OGD/R). The effects of these molecules were assessed using siRNA knockdown, conditioned media assays with SY5Y neuronal cells, in vivo overexpression of ISG15, histological and functional assessments.

Both IRF7 and ISG15 were significantly upregulated after stroke. IRF7 knockdown reduced ISG15 expression, whereas ISG15 knockdown did not affect IRF7, suggesting a unidirectional regulatory relationship. Conditioned media from microglia treated with siIRF7 or siISG15 increased SY5Y cell mortality, with a stronger effect in the siISG15 group, highlighting neuroprotective role of ISG15. ISG15 knockdown also enhanced microglial migration. Conversely, microglial ISG15 overexpression in vivo promoted a shift toward reduced neuroinflammation, improved neuronal survival, and enhanced functional recovery. Mechanistically, ISG15 stabilized NLRP3 protein but more strongly decreased its mRNA stability through accelerated degradation.

These findings demonstrate that the activation of IRF7/ISG15 axis in microglia inhibits NLRP3 Expression and improve the prognosis of ischemia/reperfusion in mice, with ISG15 exerting stronger neuroprotective effects. Targeting microglial ISG15 may offer a promising therapeutic strategy for ischemic stroke.

**MAIN POINTS:**

- ISG15 shows a stronger neuroprotective effect than IRF7 in ischemic stroke.
- ISG15 stabilizes NLRP3 protein but more strongly decreases its mRNA stability.
- Microglial ISG15 overexpression in vivo improves the prognosis of ischemic stroke in mice.

## 1. INTRODUCTION

Stroke remains the leading cause of disability and mortality among cerebrovascular diseases. Based on etiology and pathophysiology, stroke is broadly categorized as ischemic stroke (IS) or hemorrhagic stroke, with IS accounting for approximately 87% of all cases(L. Zhou et al., 2023). Acute IS is characterized by a sudden disruption of cerebral blood flow, resulting in an infarcted core surrounded by the ischemic penumbra. Therapeutic strategies aimed at preserving penumbral tissue can improve neurological outcomes. Current standard treatments of acute IS include intravenous thrombolysis, endovascular thrombectomy, antithrombotic therapy, agents that enhance microcirculation, and neuroprotective compounds (Patil, Rossi, Jabrah, & Doyle, 2022). The combined use of thrombolytics and neuroprotective agents is regarded as a front-line strategy in acute IS management. Neuroprotective agents encompass a broad spectrum of compounds, including ion channel blockers (Dibas, Al-Saad, & Dibas, 2019), antioxidants (Wu et al., 2022), anti-inflammatory agents (Y. Zhao et al., 2024), neurotrophic factors (L. Li et al., 2023), glutamate receptor antagonists (Y. B. Liang et al., 2020), mitochondrial stabilizers (Ranjbar, Komaki, Fayazi, & Zarrinkalam, 2024), and neurotransmitter modulators (Jeon et al., 2024), which collectively target multiple pathophysiological pathways.

The mechanisms underlying neuronal injury in IS are multifaceted, involving energy failure, glutamate excitotoxicity, calcium overload, oxidative stress, blood–brain barrier (BBB) disruption, platelet activation, peripheral immune cell infiltration, neuroinflammation, and dysregulated autophagy (Salaudeen, Bello, Danraka, & Ammani, 2024). Among these mechanisms, neuroinflammation plays a central role in ischemia/reperfusion-induced injury. Microglia, the resident immune cells of the central nervous system, are key mediators of post-IS inflammation. Under physiological conditions, microglia constantly monitor the brain microenvironment(van Olst et al., 2025). Following ischemia or other pathological insults, microglia undergo rapid activation and adopt diverse phenotypes, as revealed by single-cell sequencing. Despite this heterogeneity, microglial states are broadly categorized into pro-inflammatory and anti-inflammatory phenotypes. Activated pro-inflammatory microglia secrete cytokines, chemokines, and other cytotoxic mediators, which amplify neuroinflammation and thereby exacerbate blood-brain barrier (BBB) disruption, white matter injury (Z. Liang et al., 2023), and cerebral edema. Conversely, anti-inflammatory microglia facilitate tissue repair through phagocytosis of cellular debris, secretion of anti-inflammatory cytokines, and production of neurotrophic factors (Z. Liang et al., 2023). It’s widely accepted that microglia function as a “double-edged sword” in ischemic injury. Therapeutic strategies aimed at suppressing pro-inflammatory polarization while enhancing protective microglial responses may mitigate secondary brain injury and improve neurological outcomes following stroke.

The pathogenesis of IS and microglial activation are regulated by multiple signaling pathways, including purinergic receptors, Toll-like receptors (TLRs), inflammasomes, nuclear factor-κB (NF-κB), and interferon regulatory factors (IRFs). IRF3 and IRF5 promote pro-inflammatory polarization of microglia, whereas IRF4 facilitates anti-inflammatory phenotypes (Lenart, Cserep, Csaszar, Posfai, & Denes, 2024; Shui et al., 2024). IRF7 has been reported to downregulate pro-inflammatory gene expression (Cohen et al., 2014; Weichert et al., 2023), while its expression is upregulated during microglial transition to a pro-inflammatory state (Tanaka, Murakami, Bando, & Yoshida, 2015). Inflammatory stimuli such as lipopolysaccharide (LPS) or CpG oligodeoxynucleotides (CpG-ODN) markedly induce IRF7 expression following stroke. This upregulation is thought to represent a cellular self-protective response to inflammatory stress. In mouse models of ischemic preconditioning, both IRF3 and IRF7 are essential for TLR-mediated neuroprotection (Stevens et al., 2011). Following axonal injury, increased expression of IRF7 and IRF9 is observed in hippocampal microglia/macrophages (Mac1/CD11b^+^), where IRF7 may contribute to preserving BBB integrity and limiting leukocyte infiltration (Khorooshi & Owens, 2010). These findings support a context-dependent neuroprotective role for IRF7. However, conflicting evidence has also been reported. For instance, at 72 hours post-ischemia/reperfusion, iCpG-ODN treatment improves neurological outcomes and attenuates the ischemia/reperfusion-induced upregulation of both NF-κB and IRF7 (Y. Zhou et al., 2017). Furthermore, IRF7 has been shown to promote microglial production of inflammatory mediators, including IL-6 and CCL2 (Das et al., 2017; Z. Li et al., 2017). Its function in microglial polarization and prognosis of stroke remains to be clarified.

In this study, we analyzed the GEO public datasets GSE171102 and GSE212732 and identified IRF7 as a transcription factor significantly upregulated in IS, showing the greatest expression change. Using the iRegulon module in Cytoscape, we predicted downstream targets of IRF7 and identified ISG15 as a key candidate. Previous studies showed that IRF7 directly regulates ISG15 in human glioblastoma (H. Yang, Lin, Ma, Baffi, & Wathelet, 2003). ISG15 is expressed in immune cells and fibroblasts (Deng et al., 2024), and according to the Human Brain Atlas, also in microglia, astrocytes, oligodendrocytes, and neurons. Studies using ISG15 knockout (ISG15⁻/⁻) mice have shown that protein modification by ISG15 exerts neuroprotective effects after cerebral ischemia (Nakka et al., 2011). In a distal ischemia preconditioning model, ISG15 mediates hepatocyte growth factor (HGF)-induced NLRP3 activation. However, evidences also exist challenging this protective role. For example, neuronal overexpression of ISG15 impairs dendritic structure and behavioral performance (Hu et al., 2019), while neuronal exosomes enriched in ISG15 promote inflammatory cytokine production in microglia (Clarkson, Grund, David, Johnson, & Howe, 2022). These findings suggest that the neuroprotective effect of ISG15 may also be highly context-dependent and cell type-specific.

Based on these findings, we hypothesize that microglial IRF7 may exert neuroprotective effects in IS via regulation of ISG15. Using RNA interference and the OGD/R model, we proved that IRF7 regulates ISG15 expression in microglia. Notably, the neuroprotective effect of ISG15 surpasses that of IRF7. In the tMCAO mouse model, overexpression of microglial ISG15 improved post-stroke outcomes, which may be related to ISG15’s regulation of mRNA stability of NLRP3 and NF-κB.

## 2. METHODS

### 2.1 Experimental animals

Male C57BL/6 mice (25-30g) were provided by Chengdu Dashuo Laboratory Animal Company. All experimental procedures were approved by the Ethics Committee of the University of Electronic Science and Technology of China and carried out in strict accordance with the National Institutes of Health Guide for the Care and Use of Laboratory Animals (8th edition, revised 2010).

### 2.2 Transit middle cerebral artery occlusion/ reperfusion (tMCAO/R) model

As described previously (Penzes et al., 2020), mice were anesthetized with 4% isoflurane in medical oxygen using a plastic mask and the right common carotid artery (CCA) was exposed by surgery. The CCA and external carotid artery (ECA) were ligated with 6-0 silk sutures. A latex-coated monofilament (Beijing Cinontech, China) was then inserted through the CCA into the internal carotid artery (ICA) to occlude the middle cerebral artery (MCA). After 60 minutes of occlusion, the filament was gently withdrawn to allow reperfusion. The incision was sutured, and the mice were returned to their home cages for postoperative recovery.

### 2.3 TTC staining

Mice were anesthetized and euthanized by cervical dislocation. Brains were immediately harvested, placed in a brain matrix, and sectioned coronally into 1-mm-thick slices. The slices were incubated in 1% triphenyltetrazolium chloride (TTC) solution (Beyotime, China) at 37°C in the dark for 30 minutes, with gentle shaking every 5 minutes. Following staining, slices were fixed in 4% paraformaldehyde (PFA) for 10 minutes, then arranged in order and photographed for analysis.

### 2.4 Behavioral tests

The differences of neuronal injury between the control group and the tMCAO group were compared using Zea Longa method (scale 0-4), a widely used method for evaluating post-stroke deficits in rodents (Penzes et al., 2020). Higher scores indicate more severe neurological impairment: 0 = no deficit; 1 = incomplete extension of the contralateral forelimb; 2 = circling to the hemiplegic side while walking; 3 = leaning toward the hemiplegic side; 4 = inability to walk spontaneously with reduced consciousness. The modified DeSimoni neuroscores were used to compare the neuronal injury between the NC group and the OE-ISG15 group 24 hours after reperfusion as described previously(De Simoni et al., 2003; Orsini et al., 2012). General health and focal neurological deficits were scored separately (grade criteria are listed in supplementary Table 1 and supplementary Table 2), including body symmetry, coordination, grip strength, grooming, alertness, exploratory activity, and postural control. The maximum score is 41, with higher scores indicating greater neurological impairment.

Rotarod test was also used to evaluate motor coordination and fatigue resistance of mice. Animals were placed on an accelerating rotarod (acceleration: 20 r/min²; maximum duration: 8 minutes), and both the latency to fall and falling speed were recorded. Prior to tMCAO, mice were trained on the rotarod three times daily for three consecutive days, with 30-minute intervals between sessions. The performance in the final training session was used as a baseline. The test was conducted 24 hours post-ischemia/reperfusion.

Vestibulomotor function was assessed using the beam balance test. Mice were pre-trained to cross a narrow beam (30 × 1.3 cm²) for three days before surgery. Test was performed 24 hours after ischemia/reperfusion. The time taken to cross the balance beam and vestibulomotor reflex performance were recorded. According to a previously established scoring system(Nakka et al., 2011), performance was graded on a scale of 0 to 6: 0 = steady balance; 1 = grasps beam edge; 2 = one limb slips but maintains grip; 3 = two limbs slip or spinning occurs without falling; 4 = falls within 40-60 seconds; 5 = falls within 20-40 seconds; 6 = falls within 20 seconds. More points mean more deficits.

### 2.5 Immunofluorescence staining

Coronal brain sections (25 μm thickness) were washed three times with PBS (pH 7.4), followed by incubation with 0.5% NaBH₄ for 30 minutes. Antigen retrieval was performed by incubating the sections in sodium citrate buffer (pH 6.0) for 30 minutes. Subsequently, sections were blocked in blocking buffer (0.5% Triton X-100, 3% goat serum, and 3% BSA) on a shaker at room temperature for 4 hours. After blocking, brain sections were incubated overnight at 4 °C with primary antibodies diluted in blocking buffer. The next day, sections were washed and incubated for 1 hour at room temperature with species-specific secondary antibodies diluted in blocking buffer. After PBS washes for three times and desalination with deionized water, sections were mounted using a DAPI-containing antifade mounting medium and stored in dark until imaging. The primary and secondary antibodies used in this part were listed in supplementary Table 3. Microglial morphology was analyzed using the AnalyzeSkeleton plugin in ImageJ software.

### 2.6 Western Blotting

Tissue or cell samples were lysed in high-efficiency RIPA buffer containing phenylmethylsulfonyl fluoride (PMSF) and protease/phosphatase inhibitors (Solarbio, China) on ice for 10 minutes. Lysates were centrifuged at 12,000 rpm for 30 minutes at 4 °C, and the supernatants were collected. Protein concentration was determined using a BCA protein assay kit (Solarbio). After normalization of protein concentrations, samples were denatured by heating at 98 °C for 10 minutes, cooled on ice, and stored at -20 °C until use. Protein samples were separated on 10% SDS-PAGE gels prepared using a gel preparation kit (Yamei Biotech, China). Gels were placed in an electrophoresis chamber with running buffer (containing Tris 25 mM, Glycine 192 mM, SDS 0.1% w/v), and electrophoresis was carried out at 120 V for 50 minutes. Following electrophoresis, proteins were transferred to PVDF membranes using a pre-cooled transfer buffer at 105 V for 20-60 minutes, depending on molecular weight. Membranes were blocked in 5% non-fat milk (prepared in Tris-buffered saline with Tween 20 (TBST)) at room temperature for 1 hour, followed by incubation with primary antibodies (diluted in 5% milk) overnight at 4 °C on a shaker. TBST contains 20 mM Tris-HCl, 150 mM NaCl, and 0.1% (v/v) Tween 20 and the pH was adjusted to 7.4. The membranes were then washed three times with TBST and incubated with HRP-conjugated secondary antibodies (diluted in TBST) for 1 hour at room temperature. After three additional washes with TBST, immunoreactive bands were visualized using enhanced chemiluminescence (ECL, Thermo Scientific, USA), and band intensities were quantified using ImageJ software.

### 2.7 RT-qPCR

Total RNA was extracted from the injured brain hemisphere of mice and cultured cells with TRIzol reagent (Yeasen, Shanghai, China). Equal amounts of RNA samples were reverse transcribed into cDNA according to the instructions of the HiFiScript gDNA Removal cDNA Synthesis Kit (CWBIO, China). Real-time qPCR was performed using SuperStar Universal SYBR Master Mix (CWBIO). Values were averaged for triplicate assays, and the relative amounts of mRNA were normalized to GAPDH using 2^-ΔΔCt^ method. Primers used in this study are listed in supplementary Table 4.

### 2.8 Cell culture

Cryopreserved cells were rapidly thawed in a 37°C water bath and immediately washed with pre-warmed Dulbecco’s Modified Eagle’s Medium (DMEM, Gibco) supplemented with 2% fetal bovine serum (FBS, Gibco). After centrifugation and removal of the supernatant, cells were resuspended in complete culture medium consisting of DMEM with 10% FBS and 1% PS (Penicillin-Streptomycin, Gibco), seeded into culture dishes, and incubated at 37°C in a humidified atmosphere containing 5% CO₂.

### 2.9 Oxygen and glucose deprivation (OGD) of cells

The culture medium of BV2 cells grown in 12-well plates was replaced with glucose-free DMEM (Pricella, Wuhan, China). The plates were then placed in a hypoxia chamber (MIC-101, Embrient Inc.) and incubated at 37 °C under a gas mixture of 94% N₂, 5% CO₂, and 1% O₂ for 2 hours to induce OGD. After OGD treatment, the medium was replaced with normal culture medium (NCM, DMEM supplemented with 10% FBS and 1% PS), and cells were returned to a normoxic incubator for an additional 24 hours before collection.

### 2.10 RNA interference

BV2 cells were transfected with 100 nM of either negative control siRNA (NC) or siRNAs targeting IRF7 (siIRF7) and ISG15 (siISG15) (Tsingke Biotechnology Co., Ltd., China) using Lipofectamine™ 2000 (ThermoFisher Scientific, USA) according to the manufacturer’s instructions. The siRNA target sequences were obtained from the MERCK database (listed in supplementary Table 4). Cells were harvested 48 hours post-transfection for mRNA analysis and 72 hours post-transfection for protein detection.

### 2.11 Calcein-AM/ propidium iodide (PI) staining

SY5Y cells were seeded into 96-well plates. BV2 cellss were transfected with NC or siRNA and subjected to OGD. After 24 hours of recovery under normal culture conditions, the conditioned medium of microglia was collected and used to treat SH-SY5Y cells for 24 hours. SY5Y cells were rinsed in PBS and dyed with 2 μM Calcein-AM and 4.5 μM PI provided by the assay kit (Beyotime, China) at 37 °C for 30 min. After wash the cells with PBS gently, an inverted fluorescence microscope (ZEISS) was applied to take the images. The cell mortality in each field of view (%) = (the number of PI^+^ cells/total number of cells) ×100%.

### 2.12 Cell migration assay

BV2 cells were seeded at low density in 12-well plates with NCM and incubated under normoxia for 12 h. Then cells were transfected with siRNA and subjected to OGD by culturing in glucose-free medium under hypoxia (1% O₂, 5% CO₂, 94% N₂) for 2 h. After OGD, the medium was replaced with NCM for recovery. Upon reaching ∼100% confluence, a scratch was made using a 200 μL pipette tip. Wells were gently washed 3–5 times with PBS to remove debris, and images of the scratch were captured immediately (0 h). Medium was replaced with low-serum DMEM (2% FBS), and cells were incubated at 37 °C, 5% CO₂. Scratch areas were imaged at 12 and 24 h post-scratch to evaluate cell migration. The scratch width was measured using ImageJ software, and the cell migration rate (%) was calculated as follows:

Migration rate= (Initial width−Final width)/ Initial width×100%.

### 2.13 Phagocytic activity assay

BV2 cells were seeded at low density in 24-well plates and incubated under normoxic conditions for 12 h. Cells were then transfected with siRNA and subjected to OGD for 2 h, followed by restoration to normal culture conditions. After 22 h, fluorescent latex beads (1.0 μm; Sigma) diluted 1:1000 in culture medium were added and incubated for 2 h under standard conditions. The medium was then removed, and cells were gently washed three times with ice-cold PBS to remove non-phagocytosed beads. Cells were fixed with 4% paraformaldehyde at room temperature for 15 min and washed with PBS. Random fields were imaged using a fluorescence microscope, and phagocytic capacity was quantified as the mean number of internalized fluorescent beads per cell.

### 2.14 Virus vector construction and production

The pAAV-IBA1-ISG15 construct was generated in-house using Gibson Assembly. The pAAV promoter was replaced with the IBA1 promoter sequence, and the coding sequence of ISG15 (retrieved from NCBI and UCSC databases) was inserted. Cloning was performed with the Trelief® SoSoo Cloning Kit (TSINGKE), and restriction enzymes were obtained from NEB (New England BioLabs, USA). The pAAV-MG1.2 (#184541) and pHelper plasmids were purchased from Addgene. All plasmids were verified by Sanger sequencing (Sangon Biotech, China). AAV particles were produced by triple-plasmid transfection of HEK293 cells using Lipofectamine™ 2000 (Invitrogen, Thermo Fisher Scientific, USA). Seventy-two hours post-transfection, cells were harvested, and AAV was purified using AAVpro® Purification Solution (Takara, Japan). AAV-IBA1-EGFP was used as the control virus. Viral suspensions (2 μL) were stereotactically injected into the lateral ventricle (coordinates: AP -0.7 mm, ML -1.3 mm, DV -1.5 mm). tMCAO surgery was performed six weeks after AAV injection.

The lentiviral vector pLV-IRF7 was constructed from the laboratory vector LV-2.0-hSyn::HRasG12V-IRES-mCherry by replacing the original promoter with CMV and inserting the IRF7 coding sequence via AgeI and XhoI sites using Gibson Assembly. Both sequences were obtained from the NCBI database. The construct was confirmed by Sanger sequencing (Sangon Biotech) and subsequently transfected into BV2 cells. Cells were collected 48 h post-transfection for mRNA expression analysis.

### 2.15 RNA sequencing and data analysis

BV2 cells were pretreated with siISG15 or negative control (NC) and subjected to OGD for 2 h, followed by 24 h of normal culture. Total RNA was extracted using the NEBNext Ultra RNA Library Prep Kit (Illumina), and libraries were constructed according to the manufacturer’s instructions. Sequencing was performed on an Illumina HiSeq platform (Wuhan Kangce Technology Co., Ltd.; project number: KC2024-BR-1113-4546).

Raw reads were processed with Fastp (v0.23.0) to remove adapters, low-quality reads, and poly-N sequences. Clean paired-end reads were aligned to the mouse reference genome (GRCm39), and transcript abundance was calculated as counts per million. Differentially expressed genes (DEGs) between groups were identified using DESeq2 (v4.4.1), with thresholds of p < 0.05 and |log2 fold change| > 1. Gene Ontology (GO) and Kyoto Encyclopedia of Genes and Genomes (KEGG) enrichment analyses were performed with the ClusterProfiler and org.Mm.eg.db packages, covering biological process, molecular function, and cellular component categories. Gene set enrichment analysis (GSEA) was also conducted using ClusterProfiler, and pathways with adjusted p < 0.05 were considered significantly enriched.

Public datasets from GEO (GSE212732 and GSE212732) were analyzed for validation. For dataset GSE212732, rat gene symbols were converted to mouse homologs using the homologene package (v4.4.1). DEGs were identified using the same criteria as above. Protein-protein interaction (PPI) networks of upregulated DEGs were constructed using the STRING database and analyzed in Cytoscape (v3.8.2) with the CytoHubba plugin to identify hub genes. Transcription factors regulating hub genes were predicted using CHEA3, and their target genes were identified with the iRegulon module in Cytoscape.

### 2.16 Statistical analysis

Data are expressed as mean ± standard deviation (SD). Normality of data distribution was assessed using GraphPad Prism (version 8). For comparisons between two groups, an unpaired two-tailed Student’s t-test was used for normally distributed data, and a Mann-Whitney U test for non-normal data. A p-value < 0.05 was considered statistically significant.

## 3. RESULTS

### 3.1 Analysis of microglia datasets related to ischemic stroke

To identify key genes associated with IS in microglia, we analyzed the GEO dataset GSE171102. A total of 1285 DEGs were identified, including 666 upregulated and 619 downregulated genes (supplementary Figure S1b). Upregulated DEGs were enriched in biological processes such as cellular response to interferon and external stimuli, regulation of innate immunity, cytokine-mediated signaling, and chemotaxis (supplementary Figure S1a). Their gene products predominantly localized to receptor complexes, membrane structures, and the extracellular matrix (supplementary Figure S1c). KEGG analysis revealed that upregulated DEGs were mainly associated with inflammation, phagocytosis, calcium signaling, cytokine-cytokine receptor interaction pathways, and cell adhesion molecules (supplementary Figure S1d). Additionally, several antiviral signaling pathways were activated following ischemia-reperfusion. These findings align with reported pathological features of ischemia-reperfusion injury, including neuroinflammation, calcium overload, neuronal injury, complex cellular interactions for injury repair, restricted spread of inflammation, enhanced leukocyte adhesion and infiltration, microglial migration, and microglial phagocytosis of injured neurons.

The downregulated DEGs were primarily associated with biological processes including axonal myelination, wound healing regulation, homeostasis, tissue regeneration, and calcium transmembrane transport (supplementary Figure S2a). These findings align with well-known ischemia-reperfusion pathologies such as neuronal death, axonal demyelination, impaired calcium transport, disrupted nervous system homeostasis, and hindered neural regeneration. The proteins of downregulated DEGs were mainly localized to the extracellular matrix, myelin, cell membrane, and synapses (supplementary Figure S2b). KEGG analysis indicated that downregulation of cytoskeleton-related signaling pathways contributes significantly to ischemia-reperfusion-induced neuronal injury (supplementary Figure S2c).

To explore ischemia-associated alterations in gene expression at the tissue level, we further analyzed the GEO dataset GSE212732, comparing rat cortical tissue samples between the sham group and the MCAO 12-hour group. A total of 954 DEGs were identified, including 275 upregulated and 597 downregulated genes. GO biological process analysis indicated that the upregulated DEGs were predominantly associated with wound healing, hypoxia response, immune cell chemotaxis and migration, apoptosis, and responses to inflammatory stimuli and pro-inflammatory cytokines (supplementary Figure S3a). These genes were involved in the signaling pathways related to inflammation, chemotaxis of peripheral immune cells, lipid metabolism, and apoptosis (supplementary Figure S3b). GO biological process of the downregulated DEGs were predominantly associated with axons, cognition, membrane potential regulation, synaptic plasticity, neurotransmitter synthesis, transport, secretion, and regulation (supplementary Figure S4a). These downregulated DEGs were involved in the signaling pathways related to glycolipid metabolism and function of glutamatergic synapse (supplementary Figure S4b).

Microglial (GSE171102) and cortical (GSE212732) datasets both revealed significant activation of inflammation-related signaling pathways during ischemia-reperfusion, highlighting the critical role of microglial inflammation and cell interaction in tissue injury. To identify DEGs shared by microglia and cortex, rat cortical DEGs (GSE212732) were converted to mouse homologs using the homologene R package. Intersecting the DEGs from GSE212732 and GSE171102 identified 31 commonly upregulated and 39 commonly downregulated genes. The commonly upregulated genes were enriched in signaling pathways consistent with those in the cortical dataset, whereas no pathways showed significantly enriched among the commonly downregulated genes under the selected criteria. PPI analysis of the 666 upregulated DEGs from the microglial GSE171102 dataset was performed using the STRING database (confidence score ≥ 0.7). The CytoHubba plugin in Cytoscape 3.8.2 identified 11 hub genes (including Usp18, Ifi44, Ifit1, Ifih1, Ifit3, Ddx58, Isg15, Rsad2, Rtp4, Irf7, Oasl2) (Figure 1a). Using the CHEA3 web tool, we predicted 1294 transcription factors regulating these genes, among which 34 were also upregulated DEGs (Figure 1b). Among these, IRF7 exhibited the highest fold change (Figure 1c) and was mainly involved in inflammation- and antiviral-related pathways (Figure 1d). Target genes of IRF7 were predicted using the iRegulon module in Cytoscape, identifying ISG15 as one of its targets (Figure 1e).

**Figure 1.**
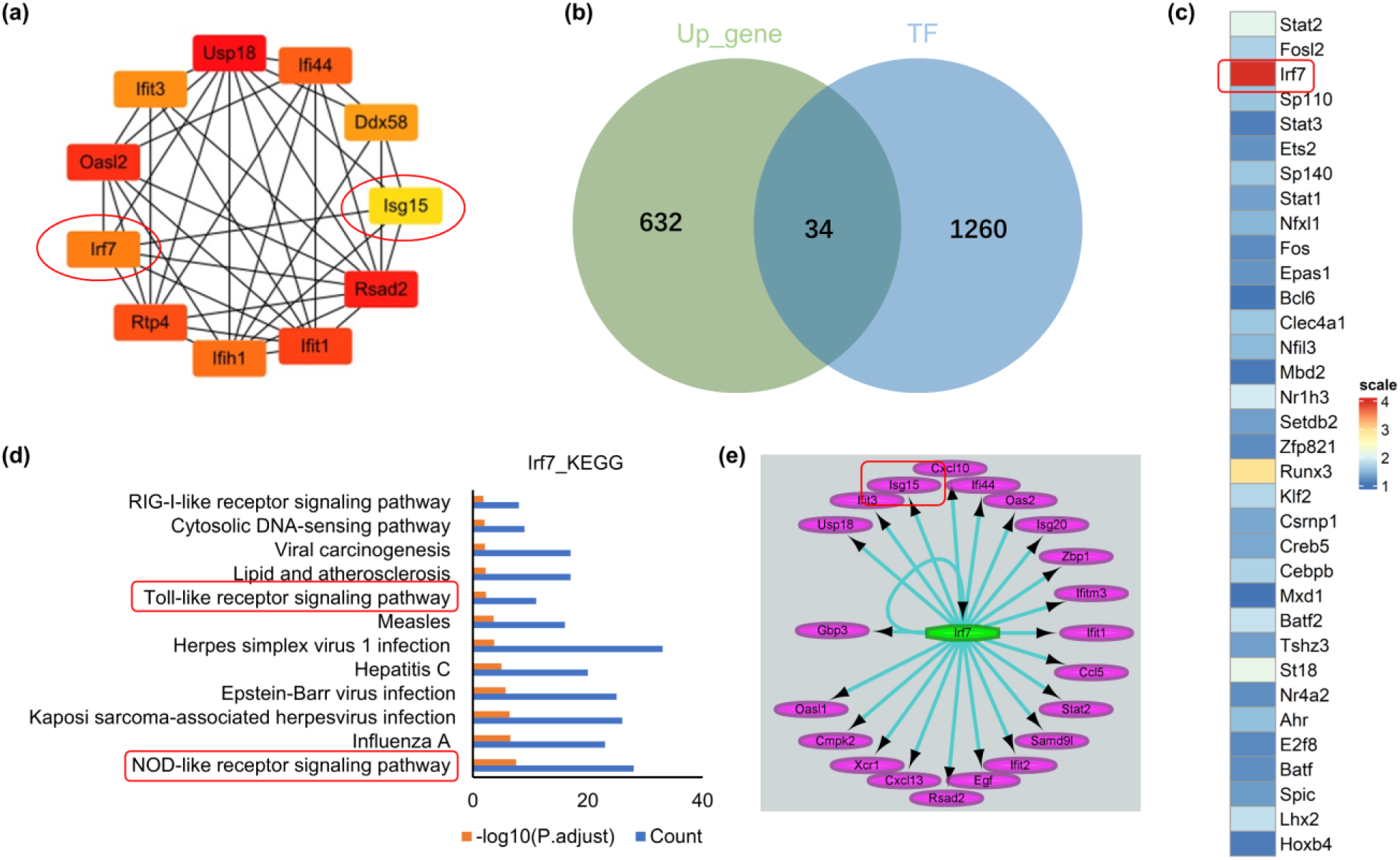
Hub gene analysis for GSE171102. (a) The protein interaction network of 11 Hub genes. (b) Among the 1294 transcription factors predicted to regulate the 666 upregulated DEGs, 34 were also identified as upregulated DEGs. (c) Among the 34 upregulated transcription factors, IRF7 exhibited the highest fold change. (d) Analysis of IRF7-related signaling pathways shows that IRF7 is mainly involved in inflammation- and antiviral-related pathways. (e) Target genes of IRF7 are predicted using the iRegulon module in Cytoscape, identifying ISG15 as one of its targets.

In human glioblastoma, IRF7 can directly regulates ISG15. Clinical data link ISG15 to several ischemic stroke subtypes, including cardioembolic, large artery atherosclerotic, and small vessel occlusive strokes (Chong et al., 2019). ISG15 is predominantly expressed in immune cells and fibroblasts (Kang, Kim, & Jeon, 2022), and the Human Brain Atlas shows its expression in microglia, astrocytes, oligodendrocytes, and neurons. ISG15 is a ubiquitin-like modifier that covalently conjugates to proteins via its C-terminal diglycine motif, forming reversible ISG15ylation. This modification regulates viral infection, cancer progression, hypoxic response, and exosome secretion. Studies using ISG15⁻/⁻ mice have shown that increased ISG15ylation after cerebral ischemia exerts neuroprotective effects (Nakka et al., 2011), though mechanisms remain unclear. In remote ischemic preconditioning, HGF from the liver protects against ischemic stroke by upregulating ISG15 in brain tissue, acting on microglial ISG15 to suppress NLRP3 activation (Yu et al., 2023). Based on the literatures and bioinformatics analyses, we hypothesized that microglial IRF7 takes protective effect in transient ischemic stroke by upregulating ISG15 and downregulating NLRP3.

### 3.2 IRF7 and ISG15 are significantly upregulated in the injured brain after ischemic stroke

To investigate the role of IRF7 and ISG15 in ischemic stroke, tMCAO model was established using a latex-coated filament. Cerebral blood flow of the brain was monitored by laser doppler flowmetry, revealing a marked reduction of blood flow of injured brain side following occlusion (supplementary Figure S5a). TTC staining performed 24 hours after reperfusion showed distinct infarct areas (white) on the infarcted hemisphere in tMCAO mice, while the non-infarcted brains remained uniformly red (supplementary Figure S5b). Neurological deficits were assessed after reperfusion. tMCAO mice exhibited significantly impaired locomotor performance in the rotarod test, with an average latency of 10.8 seconds compared to 102.7 seconds in sham controls (supplementary Figure S5c), and a significantly reduced fall speed (supplementary Figure S5d).

Neurological severe score ranged from 1 to 4 in the tMCAO group, whereas sham mice showed no deficits (score = 0) (supplementary Figure S5e). Results of the beam balance test were consistent with neurological severe score, further confirming locomotor impairment in the tMCAO group (supplementary Figure S5f).

mRNA and protein levels of phosphorylated IRF7 (pIRF7) and ISG15 were assessed 24 hours after tMCAO. Both pIRF7 and ISG15 were significantly upregulated in the cortex and hippocampus of the injured (right side) hemisphere (Figure 2a-f). Immunofluorescence analysis showed that IRF7 expression strongly co-localized with IBA1, suggesting that its upregulation following tMCAO is primarily restricted to microglia (Figure 2g, h). ISG15 was predominantly expressed in the peri-infarct (penumbra) region, with lower expression in the ischemic core and almost no signal in non-ischemic areas (Figure 2i). Unlike IRF7, ISG15 showed co-localization with microglia, astrocytes, and neurons (Figure 2j). To assess the impact of ischemic stroke on microglial phenotype, microglial morphology was compared between sham group and tMCAO group. In the tMCAO group, microglia exhibited significantly enlarged soma, reduced terminal endpoints, and decreased branches and maximum branch length (supplementary Figure S6). These morphological changes are indicative of activated microglial state, which is associated with enhanced migratory capacity and phagocytic activity.

**Figure 2.**
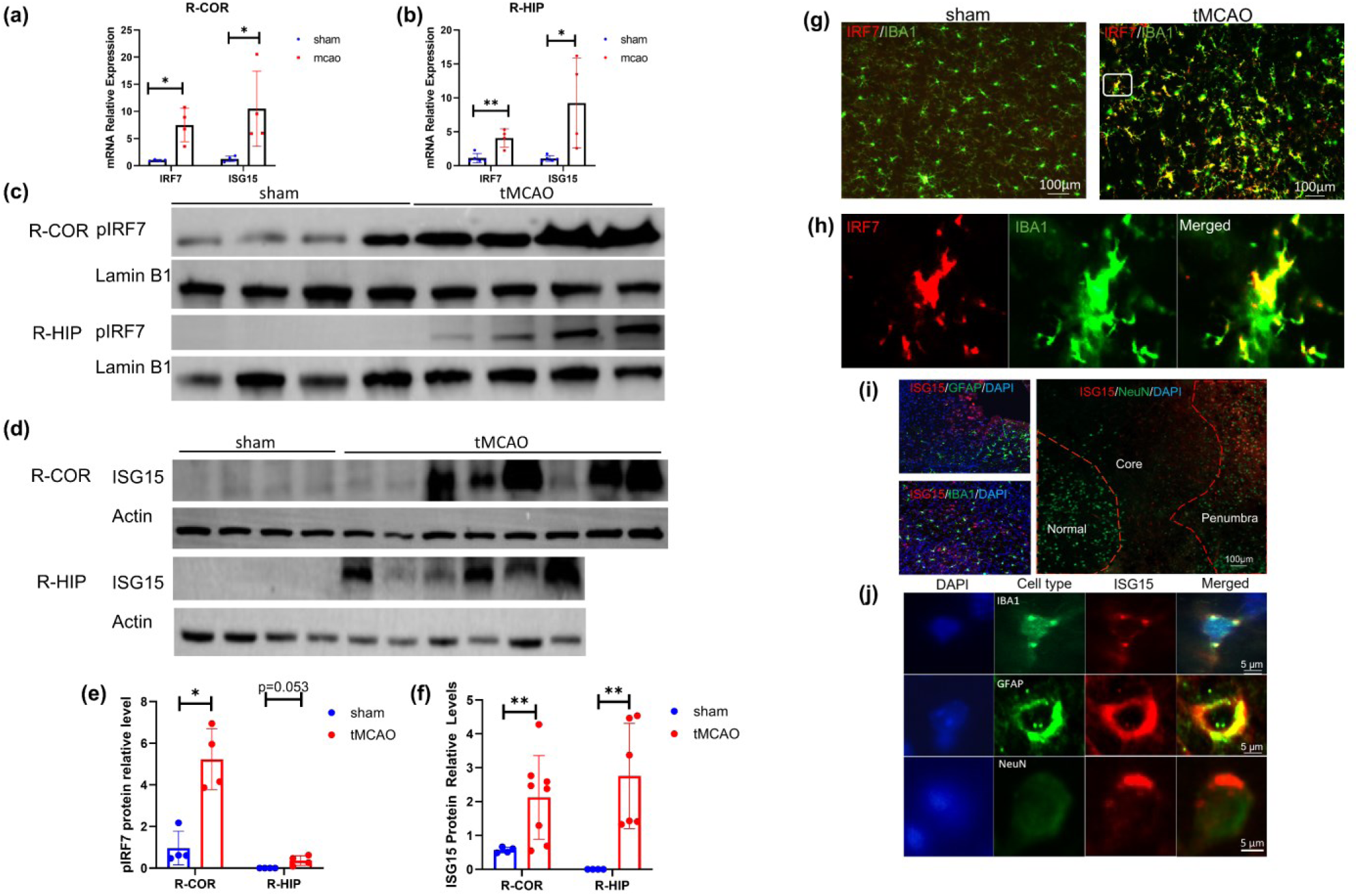
Expression and localization of pIRF7 and ISG15 mRNA and protein at 24 hours after tMCAO. (a-b) IRF7 and ISG 15 mRNA expression were significantly increased in the right cortex (R-COR) (n=4) and in the right hippocampus (R-HIP) of the injured brain (n=4). (c-d) plRF7 and ISGl5 protein expression in the R-COR and the R­ HIP of the injured brain. (e) plRF7 protein expression was increased in the R-COR and R-HIP of the tMCAO group (n = 4 for both sham and tMCAO groups). (f) ISG I5 protein expression was increased in the R-COR and R-HIP of the tMCAO group (n=4 for the sham group, n=8 for the R-COR of tMCAO group, and n=6 for the R-HIP of tMCAO group). (g, h) IRF7 expression strongly co-localized with lBAl. Localization of IRF7 in a single microglial cell (h). (i) JSG15 was predominantly expressed in the peri-infarct (penumbra) region, with lower expression in the ischemic core and almost no signal in non-ischemic areas. (i-j) ISG15 showed co-localization with microglia (marked with IBA I), astrocytes (marked with GFAP), and neurons (marked with NeuN) in the injured brain. Colocalization of ISG I5 with GFAP, IBA! and NeuN in single cells (j). Data are expressed as mean±S.D, * *P<0.05, **P<0.01*.

### 3.3 Inflammation-related molecules are upregulated in the injured brain of tMCAO mice

Transcriptomic analysis revealed that IRF7 is involved in multiple inflammation-related pathways, including the NOD-like and Toll-like receptor signaling pathways. NLRP3, GSDMD, IL-1β serve as main components of the NOD-like receptor signaling pathways. The NLRP3 inflammasome induces pyroptosis and promotes inflammatory responses by cleaving downstream GSDMD and IL-1β via caspase-1. NF-кB is involved in both pathways and is widely regarded as a key mediator of inflammation. It has been reported that that ISG15 regulates NLRP3. IRF7 may exert its regulatory effect on NOD-like receptor signaling pathway and inflammation through the modulation of ISG15. Therefore, we assessed the expression of these inflammation-related molecules. In the cortex of the injured brain (right side) of tMCAO mice, expressions of inflammation-related molecules showed an increasing trend (supplementary Figure S7a, c). In the hippocampus of injured brain, these changes were more pronounced, with inflammasome-related molecules (NLRP3, NF-κB, precursor of GSDMD (GSDMDpre), IL-1β) significantly being upregulated in the tMCAO group (supplementary Figure S7b, d).

We also examined inflammation-related molecules in the non-injured (left) hemisphere (supplementary Figure S7e-h), which may reflect the activation of anti-inflammatory mechanisms in the contralateral hemisphere. Both NLRP3 and precursor of IL-1β (IL-1βpre) were significantly downregulated in both the left hippocampus and the left cortex. GSDMD and IL-1βmat were almost undetectable in both the left cortex and hippocampus (supplementary Figure S7e, f). Collectively, these findings suggest that inflammasome signaling is robustly activated in the injured hemisphere following tMCAO, whereas the non-injured hemisphere shows the activation of anti-inflammatory response.

### 3.4 OGD/R treatment significantly increases the expression of IRF7, ISG15 and inflammation-related molecules in microglia

To further investigate the molecular mechanism of IRF7 upregulation in inflammasome activation following tMCAO, microglial cells were subjected to OGD for 2 hours, followed by 24 hours of reoxygenation to mimic the pathophysiological process. Both mRNA and protein levels of pIRF7 and ISG15 were significantly elevated after OGD/R (Figure 3a–e). In addition, expression of inflammation-related molecules was markedly increased (Figure 3f, g), consistent with in vivo findings.

**Figure 3.**
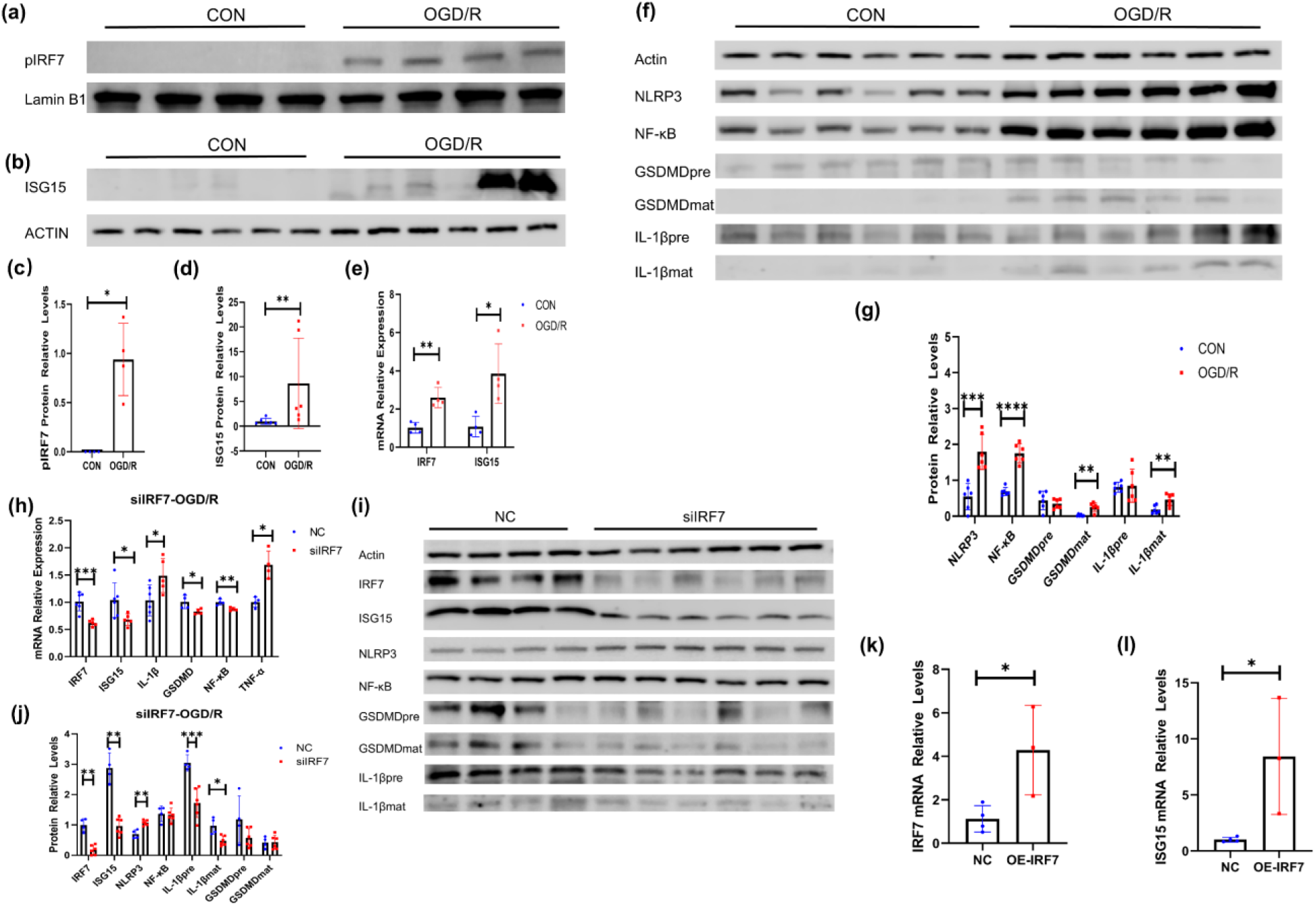
Effects of OGD/R on the expression of pIRF7, lSG 15, and inflammasome-related molecules in microglia. (a-b) Protein bands of pIRF7 and ISG 15. (c-d) pIRF7 (n=4 for each group) and ISGI 5 (n=6 for each group) protein expression were significantly increased in the OGD/R group. (e) IRF7 and ISGI5 mRNA expression (n=4 for each group) were significantly increased in the OGD/R group. (f) Protein bands of inflammasome-related molecules. (g) Expression of inflammation-related molecules was markedly increased in the OGD/R group (n=6 for each group). (h) Knockdown of IRF7 significantly reduced the mRNA levels of IRF7, ISG15, GSDMD and NF-KB, while significantly increased mRNA levels of IL-IP and TNF-a; for the assays of IRF7, ISGI5, and IL-IP, n = 6 for the NC (negative control) group and n = 5 for the silRF7 group, while for the assays of GSDMD, NF-KB, and INF-a, n = 4 for each group. (i) Protein bands of inflammation­ related molecules. U) Knockdown oflRF7 significantly reduced the protein expression oflRF7, JSG15 and IL-Ip, while significantly increasing the protein level ofNLRP3 in the silRF7 group (n=4 for the NC group, n=6 for the silRF7 group). (k) Transfection of cells with the IRF7 overexpression vector significantly increased IRF7 mRNA expression, demonstrating the effectiveness of IRF7 overexpression (n=4 for the NC group, n=3 for the OE-IRF7 group). (I) IRF7 overexpression significantly increased TSG I5 mRNA level (n=4 for the NC group, n=3 for the OE-TRF7 group). Data are expressed as mean±S.D, *\*P<0.05,* **P<0.01, ***P<0.001, ****P<0.0001 . CON: blank control.

### 3.5 Microglial IRF7 exerts a protective role in OGD/R

OGD/R treatment was shown to upregulate both IRF7 and inflammation-related molecules. To further clarify the role of IRF7 in the inflammatory response, microglial cells were pretreated with siIRF7 or a negative control (NC) prior to OGD/R. Knockdown of IRF7 significantly reduced both mRNA and protein levels of ISG15 (Figure 3h-j), whereas overexpression of IRF7 significantly increased ISG15 mRNA expression (Figure 3k-l), supporting the role of IRF7 as a regulator of ISG15 (Schematic diagrams of the overexpression plasmid of IRF7 and negative control plasmid (NC) were shown in supplementary Figure S8).

In addition, siIRF7 pretreatment significantly reduced NF-κB mRNA levels; however, its protein expression remained unchanged following OGD/R. Interestingly, NLRP3 protein expression was markedly elevated in the siIRF7 group compared with the NC group. While the mRNA level of IL-1β increased after siIRF7 treatment, its protein expression was reduced. Although the mRNA level of GSDMD was significantly decreased by siIRF7 pretreatment, its protein expression remained unchanged by siIRF7 (Figure 3h–j). Given these complex effects of siIRF7 on the expression of inflammation-related molecules, we further treated SY5Y cells for 24 hours with conditioned medium derived from microglia of the siIRF7 and NC groups to clarify the role of microglial IRF7 in OGD/R. The death rate of SY5Y cells increased from 10% in the NC group to 20% in the siIRF7 group, suggesting that IRF7 upregulation following OGD/R may exert a neuroprotective effect (Figure 4d, e).

**Figure 4.**
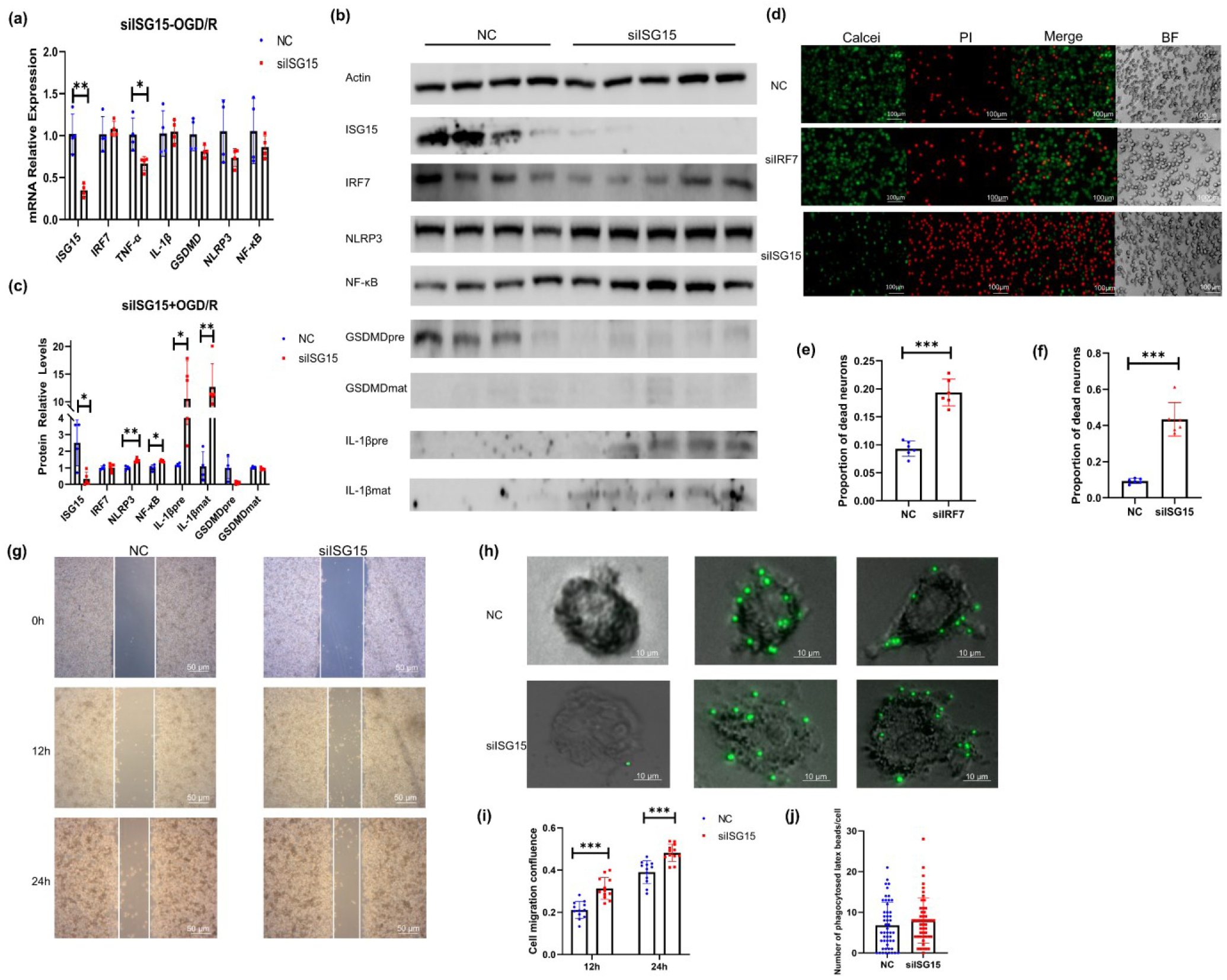
The effect of silSG15 on physiological characteristics of microglia after OGD/R. (a) Knockdown of ISG15 using siRNA (silSG 15) significantly reduced the mRNA levels of ISG15 and TNF-a. (b) Protein bands of inflammation-related molecules. (c) silSG 15 significantly reduced the protein expression of ISG15, while significantly increasing the protein levels of NLRP3, NF-KB, and IL-Ib(n=4 for the NC group, n=5 for the silSG 15 group). (d) Calcein-AM/PI staining of SY5Y cells after co-culture with conditioned medium from microglia subjected to si!RF7, silSG15, and OGD/R treatment. (e-l) The death rate of SY5Y cells increased in both si1RF7 and silSGl5 groups (n=6 for each group). (g) Representative images showing the migration of microglia at different time points. (h) Representative images showing microglial phagocytosis of beads. (i) silSG 15 significantly enhanced microglial migration at 12 and 24 hours post-OGD/R (n=l 2 for each group). (j) Microglial phagocytic capacity remained unchanged in silSG15 group (n=50 cells for each group). Data are expressed as mean±S.D, *\*P<0.05,* ** *P<O.O*I,*\*\*\*P<0.00* I.

### 3.6 ISG15 plays a stronger protective role in OGD/R

To further investigate the role of ISG15 in microglial inflammation, microglial cells were pretreated with siISG15 followed by OGD/R. Knockdown of ISG15 significantly reduced mRNA levels of both ISG15 and TNF-α, while mRNA levels of other inflammatory molecules remained unchanged (Figure 4a). In contrast, protein expression of NLRP3, NF-κB, IL-1βpre and IL-1βmat were significantly increased (Figure 4b, c). These findings suggest that siISG15 exacerbates OGD/R-induced inflammation in microglia. Notably, siISG15 had no effect on either mRNA or protein levels of IRF7, indicating that ISG15 does not regulate IRF7 in a feedback manner.

Similarly, conditioned medium from microglial cells pretreated with siISG15 and subjected to OGD/R was applied to SY5Y cells for 24 hours. The mortality of SY5Y cells increased from 10% in the NC group to 40% in the siISG15 group (Figure 4d, f), indicating that ISG15 upregulation following OGD/R exerts a robust neuroprotective effect-potentially stronger than that of IRF7.

### 3.7 siISG15 treatment increases microglial migration without significantly altering their phagocytosis

The inflammatory environment following stroke promotes microglial migration toward the ischemic core and penumbra. To assess the role of ISG15 in this process, scratch assays were performed in microglial cells transfected with siISG15 and subjected to OGD/R. Microglial migration was significantly increased at both 12 and 24 hours after OGD/R in the siISG15 group compared to NC group (Figure 4g, i). Microglia also contribute to injury response through cytokine release and phagocytosis of debris from apoptotic cells. To evaluate phagocytic capacity, yellow-green fluorescent latex beads were used to incubate with microglia for 2 hours. Microglial phagocytic capacity remained unchanged in siISG15 group (Figure 4h, j).

### 3.8 Overexpression of ISG15 in microglia of the injured brain improves the locomotor function of mice after ischemic stroke

Our previous studies have shown that the neuroprotective effect of ISG15 is more pronounced than that of IRF7, and that ISG15 is widely expressed in neurons, astrocytes, and microglia. However, overexpression of ISG15 in neurons has been reported to induce dendritic damage and behavioral deficits (Hu et al., 2019), and neuronal exosomes enriched in ISG15 have been shown to elevate inflammatory cytokine expression in microglia (Clarkson et al., 2022). These findings suggest that cell type-specific regulation of ISG15 may offer a safer and more effective therapeutic strategy.

Forty-five days after AAV-mediated overexpression of ISG15 via injection into the right lateral ventricle, mice were subjected to tMCAO surgery. To establish baseline motor performance, rotarod training was performed three times daily for three consecutive days prior to surgery, with no significant differences observed between the NC and OE-ISG15 groups (Figure 5a–c). At 24 hours after tMCAO and reperfusion, mice overexpressing ISG15 showed markedly improved locomotor function, as indicated by a significant increase in latency to fall from the rotarod in the OE-ISG15 group (Figure 5d). In addition, both general neurological deficit scores and focal neurological severity scores were significantly lower in the OE-ISG15 group (Figure 5e). Consistently, infarct volume in the injured hemisphere was substantially reduced (Figure 5f, g). The results indicate the protective effect of up-regulated microglia ISG15 in mice brain after ischemic stroke.

**Figure 5.**
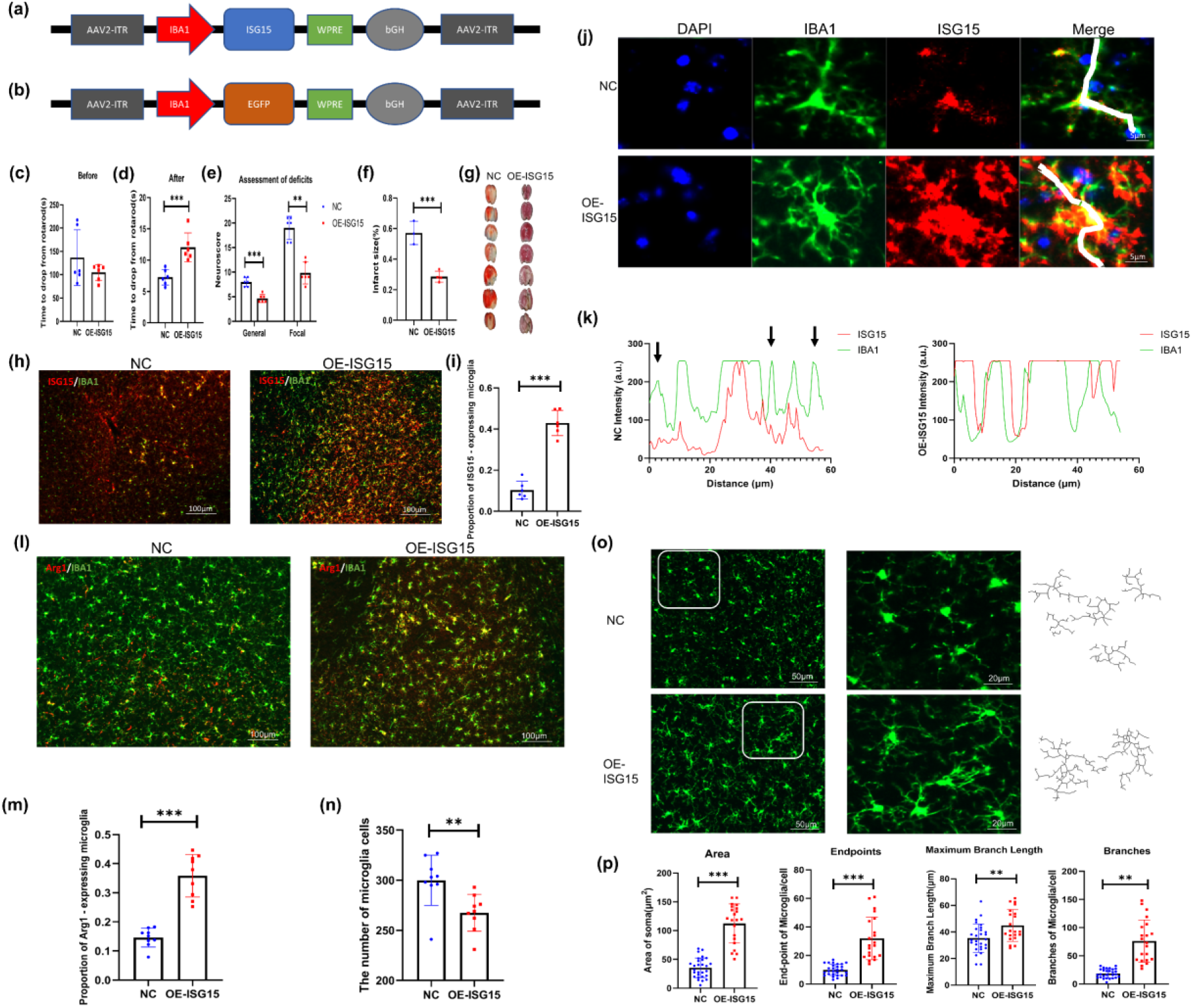
Effects of ISG15 overexpression on neurological injury after stroke, as well as on microglial density and phenotype. (a-b) Schematic diagrams of the AAV vector for ISGl5 overexpression in microglia and the corresponding negative control vector. (c) There was no significant difference between the NC and OE-ISG I5 groups in the latency of mice to drop from the rotarod before tMCAO (n=6 for each group). (d) At 24 hours after tMCAO and reperfusion, mice overexpressing ISG15 showed marked increase in the latency to fall from the rotarod (n=6 for each group). (e) General and focal neurological injury scores significantly decreased in mice overexpressing !SG15 (n=6 for each group). (f) Infarct area significantly decreased in mice overexpressing ISG15 (n=3 for NC group, n=4 for OE-ISG15 group). (g) TIC staining. (h) Immunofluorescence analysis confirmed robust upregulation of ISG15 in IBA I+ microglia in the OE-ISG15 group. (i) The proportion of ISG I 5+ microglia significantly increased in the OE-ISGl5 group (n=6 for each group). (j) Co-localization studies showed that in the OE-ISGI5 group, ISGI5 was present in both microglial soma and processes, extending beyond the IBA! signal into the extracellular space. In contrast, in the NC group, ISGl5 localization was largely restricted to the soma with much fewer process expression. (k) Fluorescence colocalization analysis along the white line in (j), with black arrows pointing to regions lacking expression of ISGl5. (I) lmmunofluorescence colocalization of Argl and IBAI on the injured side. (m) There was a significant increase in the proportion of ArgI+ microglia in the OE-ISGl5 group (n=9 for each group). (n) There was a significant reduction in the number of microglial cells in the OE-ISG15 group (n=9 for each group). (o) The lmmunofluorescence and cytoskeleton images of the two groups. (p) Morphological analysis demonstrated that microglia in the OE-ISG I5 group (n=20) exhibited larger somata, greater numbers of tenninal endpoints and branches, and a greater maximum branch length compared with the NC group (n=28). Data are expressed as mean±S.D, *\*\*P<0.01, ***P<0.001*.

Immunofluorescence analysis confirmed robust upregulation of ISG15 in IBA1^+^ microglia (Figure 5h), with the proportion of ISG15^+^ microglia increasing from 10% in NC to 40% in the OE-ISG15 group (Figure 5i). Co-localization studies showed that in the OE-ISG15 group, ISG15 was present in both microglial soma and processes, extending beyond the IBA1 signal into the extracellular space. In contrast, in the NC group, ISG15 localization was largely restricted to the soma with much fewer process expression (Figure 5j, k, black arrows pointing to regions lacking expression of ISG15).

### 3.9 The overexpression of microglial ISG15 in vivo alters the phenotype and density of microglia

We further investigated whether in vivo overexpression of ISG15 altered the microglial phenotype. Immunofluorescence analysis revealed a significant increase in the proportion of Arg1⁺ microglia (Figure 5l, m), accompanied by a reduction in the number of microglial cells in the OE-ISG15 group (Figure 5n). Morphological analysis demonstrated that microglia in the OE-ISG15 group exhibited larger somata, greater numbers of terminal endpoints and branches, and a greater maximum branch length compared with the NC group (Figure 5o, p), indicating a shift toward a more ramified, anti-inflammatory phenotype.

### 3.10 The overexpression of microglial ISG15 in vivo enhances neuronal survival and alleviates neuroinflammation

To evaluate the effect of microglial ISG15 overexpression on neuronal survival, neurons were labeled with NeuN, and the survival ratio was calculated by dividing the number of NeuN⁺ cells in the ischemic hemisphere by that in the corresponding non-ischemic region. Overexpression of ISG15 significantly improved neuronal survival in the ischemic hemisphere (Figure 6a–c), as evident from the immunofluorescence staining (Figure 6b). To confirm the efficiency of ISG15 overexpression mediated by the AAV vector, the vector was injected into the lateral ventricle on the injured side (right hemisphere), and 45 days later, tMCAO was performed. Protein expression of ISG15 was assessed in the cortex and hippocampus 24 hours after reperfusion. ISG15 expression was elevated by approximately 10-fold in the right cortex and 7-fold in the right hippocampus in the OE-ISG15 group compared with the NC group (Figure 6d).

**Figure 6.**
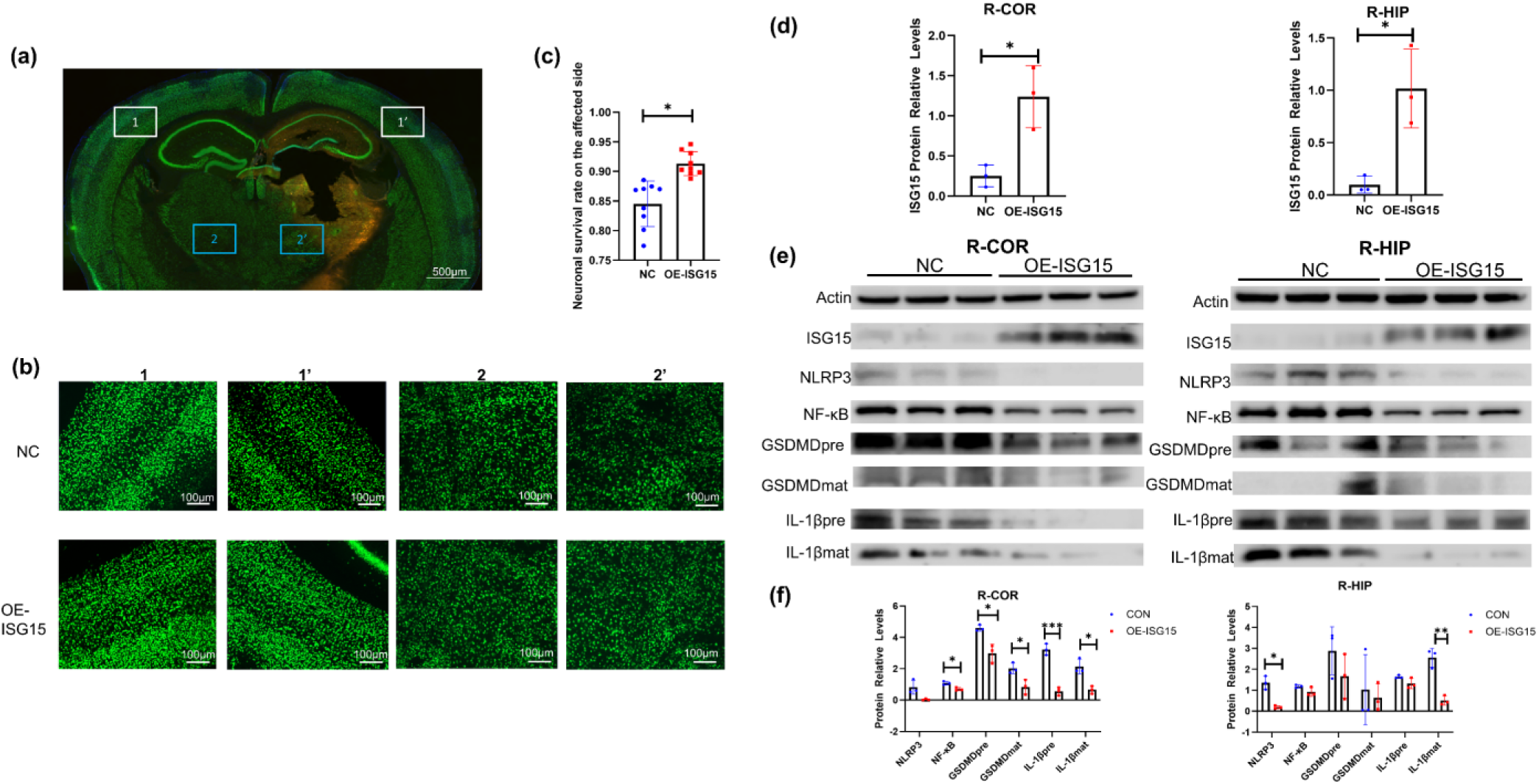
The effect of ISG15 overexpression on neuronal survival and expression of inflammation-related molecules in microglia. (a) Schematic diagram of the analyzed brain region.I and 1·, 2 and 2’ are corresponding brain regions. (b) Florescence ofNeuN in the corresponding regions. (c) Overexpression ofJSGl5 significantly improved neuronal survival in the ischemic hemisphere (three mice, with three brain slices per mouse, n=9 for each group). (d) ISGl5 expression was elevated in the right cortex (R-COR) and right hippocampus (R-HIP) of the OE-ISGl5 group compared with the NC group (n = 3 per group), with the right side representing the ischemic hemisphere. (e) Protein bands of inflammation-related molecules in the R-COR and R-HIP. (t) In the R-COR of OE­ ISG 15 mice, levels ofNLRP3, NF-KB, GSDMD, and IL-Ip were significantly reduced. In the R-HIP, NLRP3 and IL-lpmat were significantly downregulated (n=3 for each group). Data are expressed as mean±S.D, *\*P<0.05,* **P<0.01, ***P<0.001.

We next examined the expression of inflammation-related proteins. In the right cortex of OE-ISG15 mice, levels of NLRP3, NF-κB, GSDMD, and IL-1β were significantly reduced (Figure 6e, f). In the hippocampus, NLRP3 and IL-1βmat were significantly downregulated. These findings suggest that microglial ISG15 overexpression exerts a neuroprotective effect following ischemic stroke, potentially by attenuating inflammasome activation and suppressing microglial pro-inflammatory responses.

### 3.11 ISG15 enhances the protein stability of NLRP3 but decreases the mRNA stability of NLRP3

Following knockdown of ISG15 in microglia and exposure to OGD/R, no significant changes were observed in the mRNA levels of inflammation-related genes. However, protein expression of NLRP3, NF-κB, and IL-1β were significantly elevated. We hypothesized that ISG15 knockdown may enhance protein stability post-stroke. To test this hypothesis, we treated microglial cells with cycloheximide (CHX) to inhibit de novo protein synthesis and monitored the degradation of NLRP3 and NF-κB over time. At baseline (0 h), NLRP3 and NF-κB protein levels were higher in the siISG15 group than in the NC group. However, with increasing duration of CHX treatment, both proteins degraded more rapidly in the siISG15 group (Figure 7a, b). These findings suggest that ISG15 contributes to the stabilization of NLRP3 and NF-κB proteins, likely by limiting their ubiquitination, consistent with previous reports indicating that ISG15 competes for ubiquitination sites and suppresses protein ubiquitination (Y. Gao et al., 2024).

**Figure 7.**
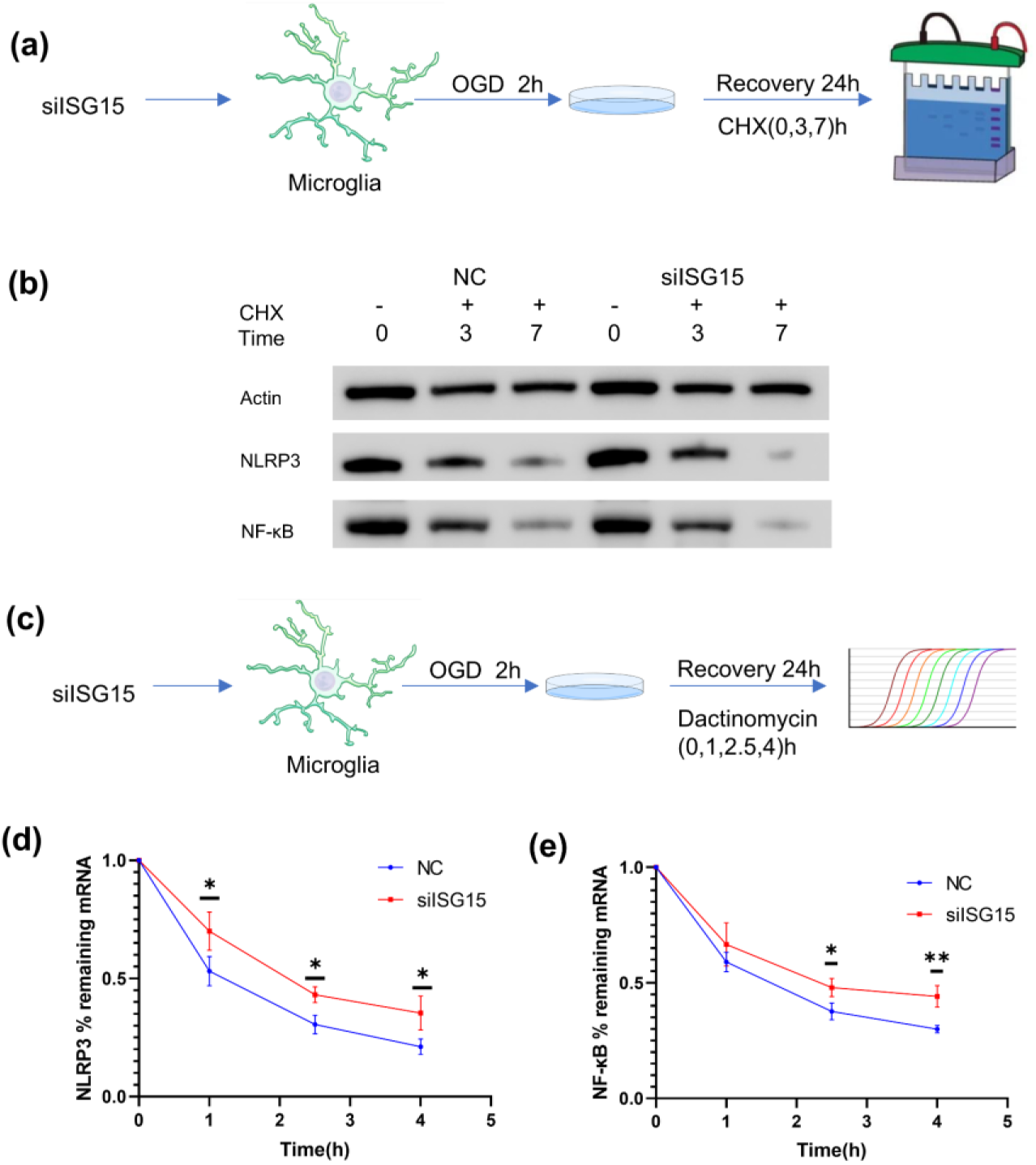
The effect of siISG15 on protein and mRNA stability in microglia after OGD/R treatment. (a) Schematic diagram of experiment. After siISG15/NC treatment, microglial cells were subjected to OGD for 2 hours, followed by 17 hours of recovery under normal culture conditions. The cells were then treated with **CHX,** and samples were collected at 0, 3 and 7 hours post-treatment for detection of NLRP3 and NF-KB protein expression by western blot. (b) Microglial cells were treated with CHX to block protein synthesis, and degradation of NLRP3 and NF-KB was monitored over time. At baseline, both proteins were higher in the siISG15 group than in the NC group and degraded more rapidly during CHX treatment.(c) Schematic diagram of experiment. After siISG15/NC treatment, microglial cells were subjected to OGD for 2 hours, followed by 20 hours of recovery under normal culture conditions. The cells were then treated with Dactinomycin, and samples were collected at 0, 1,2.5 and 4 hours post-treatment for detection of NLRP3 and NF-KB mRNA levels by RT-qPCR. (d-e) Degradation curves ofNLRP3 and NF-KB mRNA (n=3 per group). Both mRNAs were more stable in the siISG15 group at 2.5 and 4 h after actinomycin D treatment, with NLRP3 also showing increased stability at 1 h. Data are expressed as mean±S.D, *\*P<0.05, **P<0.01.* CHX: cycloheximide; Time: hour.

Our previous findings demonstrated that siISG15 significantly increased protein levels of NLRP3 and NF-κB, despite accelerating their degradation, suggesting a compensatory mechanism. Since siISG15 doesn’t change the mRNA level of NLRP3 and NF-κB, we hypothesized that ISG15 may regulate mRNA stability or translation efficiency, thereby offsetting reduced protein stability after ISG15 knockdown. To test this hypothesis, actinomycin D was used to inhibit RNA synthesis, and mRNA levels were measured at multiple time points (Figure 7c). At 0 h, baseline mRNA levels were established. In the siISG15 group, NLRP3 mRNA demonstrated significantly enhanced stability at 1, 2.5, and 4 hours following actinomycin D treatment, with its half-life prolonged from 1.1 hours in the NC group to 2.1 hours in the siISG15 group (Figure 7d). NF-кB mRNA stability was also enhanced, with no difference observed at 1 hour but significantly greater retention at 2.5 and 4 hours following actinomycin D treatment. Its half-life increased from 1.8 hours in the NC group to 2.2 hours in the siISG15 group (Figure 7e). These results suggest that knockdown of ISG15 prolongs mRNA stability of NLRP3 and NF-κB, potentially compensating for the accelerated protein degradation and contributing to increased protein expression.

### 3.12 The possible mechanisms through which ISG15 regulates the mRNA stability of NLRP3

To investigate the impact of ISG15 knockdown on microglia under ischemia-reperfusion conditions, microglial cells were transfected with siISG15, subjected to OGD/R treatment, and subsequently analyzed by RNA-seq. Comparative transcriptomic profiling revealed 591 upregulated and 186 downregulated differentially expressed genes (DEGs) in the siISG15 group relative to the negative control (NC) group (Figure 8a).

**Figure 8.**
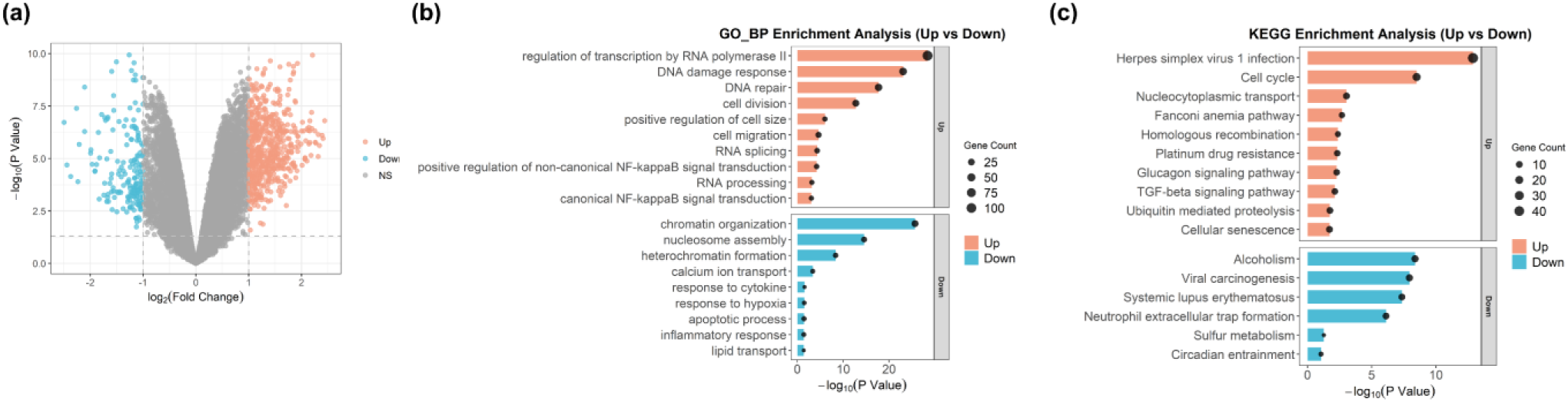
Transcriptome analysis of microglia after silSG 15 and OGD/R treatment. (a) Volcano plot of DEGs m microglia after siISG 15 and OGD/R treatment. (b) GO enrichment analysis of upregulated DEGs. The genes were mainly associated with transcriptional regulation, DNA damage and repair, cell division, cell migration, RNA splicing, and non-canonical NF-KB signaling. (c) KEGG pathway analysis of DEGs. Upregulated genes were enriched in pathways including cell cycle, glucagon signaling, TGF-P signaling, ubiquitin-mediated proteolysis, and cellular senescence, with additional enrichment of TGF-P signaling, suggesting a compensatory protective mechanism. Among downregulated pathways, neutrophil extracellular trap formation was prominent, indicating a secondary anti­ inflammatory response to lSG 15 knockdown.

GO enrichment analysis indicated that upregulated DEGs were predominantly associated with transcriptional regulation, DNA damage and repair, cell division, cell migration, RNA splicing, and non-canonical NF-κB signaling (Figure 8b). Notably, enrichment of pathways related to cell migration and NF-κB signaling was consistent with phenotypic observations, whereas the involvement of RNA splicing and processing pathways suggests a potential role of ISG15 in mRNA metabolism.

Conversely, the most significantly downregulated biological processes included chromatin organization, nucleosome assembly, heterochromatin formation, and calcium ion transport. Of particular interest, the inflammatory response category was significantly suppressed (Figure 8b). Representative genes within this category included SerpinB1A, Axl, Ccl5, IL1R2 and IL17D, which have been reported to exhibit anti-inflammatory or neuroprotective functions. For instance, SerpinB1A acts as an immune modulator that maintains antibacterial defense while attenuating tissue damage and excessive inflammation (P. Zhao, Hou, Farley, Sundrud, & Remold-O’Donnell, 2014); AXL facilitates microglial phagocytosis and clearance of Aβ plaques, and its deficiency aggravates pathology and cognitive decline (Y. Huang et al., 2021); CCL5 mitigates oxidative stress, promotes IL-10 expression, and reduces secondary damage following brain injury(Ho et al., 2021); IL1R2 functions as an IL-1 receptor antagonist, preventing IL-1 signaling and exhibiting broad anti-inflammatory activity (Peters, Joesting, & Freund, 2013); IL17D protects against inflammation in colitis models, whereas its absence exacerbates disease severity (J. Huang et al., 2021). Collectively, downregulation of these genes may contribute to the heightened inflammatory response observed following ISG15 silencing.

KEGG pathway analysis further demonstrated that upregulated DEGs were enriched in pathways such as cell cycle, glucagon signaling pathway, TGF-β signaling pathway, ubiquitin mediated proteolysis and cellular senescence, with additional enrichment of TGF-β signaling, which may represent a compensatory protective mechanism. Upregulation of ubiquitin-mediated proteolysis is consistent with the proposed role of ISG15 in ubiquitin-binding site competition. Furthermore, enrichment of the cellular senescence pathway supports the involvement of ISG15 in preserving cellular homeostasis. Among downregulated pathways, neutrophil extracellular trap formation was notable, suggesting a secondary anti-inflammatory compensatory response to ISG15 knockdown (Figure 8c).

Given that siISG15 treatment enhanced the stability of NLRP3 and NF-ΚB mRNAs, we further investigated genes associated with RNA splicing and processing. Previous studies have demonstrated that MBNL1 promotes NRF2 protein degradation by stabilizing Cul3 mRNA (Wang, Liu, & Duan, 2020), while MBNL2 enhances Bcl-2 mRNA stability through direct binding(He, Li, Chen, & Huang, 2023). Additionally, missense variants in THOC2 have been reported to impair protein stability (Kumar et al., 2020). VIRMA suppresses GSK3Β mRNA degradation (Y. Yang et al., 2024), and PTBP3 regulates the expression of the E3 ubiquitin ligase UBE4A by binding to its 3′ untranslated region, thereby preventing mRNA decay (Xie et al., 2022). These upregulated genes (MBNL1, MBNL2, THOC2, VIRMA, PTBP3) in the siISG15 group, may contribute to the increased stability of NLRP3 and NF-ΚB mRNAs through post-transcriptional regulatory mechanisms.

## 4. DISCUSSION

In this study, we provide the first evidence that microglial IRF7 exerts neuroprotective effects by modulating NLRP3 through ISG15, although its protective capacity is less pronounced compared to ISG15. Notably, microglia-specific overexpression of ISG15 markedly improved motor function and promoted tissue repair in ischemic stroke mice, primarily by attenuating neuroinflammation within the injured brain, facilitating a shift of microglial activation toward a protective phenotype, and reducing neuronal loss. Mechanistically, ISG15 enhances the stability of the NLRP3 protein while concomitantly decreasing the mRNA stability of NLRP3 and NF-κB, with a more substantial regulatory effect observed at the mRNA level. Collectively, these findings underscore the pivotal role of the IRF7–ISG15 axis in orchestrating neuroinflammatory responses following ischemia– reperfusion injury and position ISG15 as a promising therapeutic target for microglia-based interventions in ischemic stroke.

The neuroprotective role of IRF7 has been previously demonstrated. It is essential for TLR-mediated preconditioning and confers protection in models of transient cerebral ischemia. Furthermore, the TLR/IRF7/IFN signaling axis has been implicated in recovery from glutamate-induced excitotoxicity (Pappas et al., 2012). In models of cardiac arrest–induced brain injury, administration of recombinant rmMFG-E8 promotes a phenotypic shift of microglia from a pro-inflammatory to an anti-inflammatory state, accompanied by upregulation of IRF7. Notably, silencing IRF7 markedly abolishes the neuroprotective effects of rmMFG-E8 (K. Zhang et al., 2025). However, conflicting evidence exists. For instance, treatment with iCpG-ODN has been shown to improve neurological outcomes following focal cerebral ischemia-reperfusion by suppressing the upregulation of IRF7, NF-кB, and other pro-inflammatory mediators (Y. Zhou et al., 2017). Similarly, in traumatic brain injury (TBI), pharmacological blockade of hCCR2 reduces IRF7 expression and facilitates functional recovery (Somebang et al., 2021). Compounds with anti-aging properties, such as TSG, also downregulate IRF7 expression (D. Gao et al., 2023). Inhibition of the STING pathway after TBI attenuates IFN-I signaling, including IRF7 expression, thereby mitigating chronic microglial activation and cognitive decline (Packer et al., 2024). Consistent with these observations, both in vitro and in vivo studies indicate that IRF7 expression increases during the transition of microglia from an anti-inflammatory to a pro-inflammatory phenotype (Tanaka et al., 2015), underscoring its context-dependent role in neuroinflammation. In the present study, we observed a complex regulatory effect of IRF7 on inflammation-related molecules. Knockdown of IRF7 enhanced NLRP3 protein expression, possibly through downregulation of ISG15, which may stabilize NLRP3 mRNA. Although siRNA-mediated silencing of IRF7 reduced NF-кB mRNA levels (Figure 3h), NF-кB protein expression remained unchanged following OGD/R (Figure 3j). This discrepancy may result from simultaneous suppression of ISG15 at both mRNA and protein levels, which could stabilize NF-кB mRNA (Figure 7e) and lead to compensatory protein upregulation (Figure 4c). siIRF7 increased IL-1β mRNA abundance (Figure 3h), yet both precursor and active forms of IL-1β proteins were decreased (Figure 3j), potentially due to accelerated degradation of the precursor protein driven by ISG15 suppression—a hypothesis warranting further investigation. Interestingly, siIRF7 significantly reduced GSDMD mRNA expression but did not affect its protein expression (Figure 3j). This may also be related to the regulation of GSDMD mRNA stability by ISG15. Combined with the effect of conditioned medium from siIRF7-treated microglia on SY5Y cell viability, these findings suggest that IRF7 knockdown enhances the cytotoxicity of microglial secretions toward SY5Y cells under ischemia/reperfusion conditions, reinforcing the neuroprotective function of IRF7. Consistent with previous findings in human neuroblastoma cells, our results further demonstrate that IRF7 regulates ISG15 expression in microglia. Silencing either IRF7 or ISG15 markedly increases NLRP3 protein expression.

The neuroprotective role of ISG15 has been reported in previous studies. For example, the HGF/ISG15 axis has been shown to attenuate NLRP3 inflammasome activation (Yu et al., 2023). Moreover, ISG15⁻/⁻ mice exhibit increased mortality, larger infarct volumes, and poorer neurological outcomes following tMCAO compared with wild-type controls, due to the combined effects of ISG15 knockout in both the periphery and the central nervous system (Nakka et al., 2011). In the context of antiviral immunity, elevated ISG15 and PKR protein levels in the spinal cords of C57BL/6 mice restrict viral replication and confer resistance to Theiler’s murine encephalomyelitis virus (L. Li, Ulrich, Baumgartner, & Gerhauser, 2015). In addition, ISG15ylation enhances the stability of specific proteins, such as Stat1, in LPS-stimulated microglia, thereby preventing premature termination of immune responses during inflammation (Przanowski, Loska, Cysewski, Dabrowski, & Kaminska, 2018). Collectively, these findings support the notion that ISG15ylation acts as an endogenous neuroprotective mechanism. However, conflicting evidence suggests that ISG15 may also exert proinflammatory effects under certain conditions. For instance, neuronal overexpression of ISG15 has been reported to increase inflammatory cytokine production in microglia (Clarkson et al., 2022). Similarly, systemic inducible knockout of Clec16a in mice results in a neurodegenerative inflammatory phenotype resembling spinocerebellar ataxia, potentially due to upregulation of ISG15 (Hain et al., 2021). In the present study, siISG15 did not significantly alter IL-1β mRNA levels (Figure 4a), yet it markedly increased IL-1β protein expression (Figure 4c). This apparent discrepancy may be attributable to siISG15-induced upregulation of NLRP3, which facilitates inflammasome assembly and thereby promotes cleavage of the IL-1β precursor into its mature, active form. Additionally, it is conceivable that ISG15 modulates IL-1β mRNA metabolism, contributing to the observed elevation in IL-1β precursor protein levels; however, this possibility requires experimental validation. Importantly, conditioned medium from siISG15-treated microglia induced greater SY5Y cell death, likely attributable to elevated extracellular release of mature IL-1β. These findings suggest that microglial ISG15 exerts robust neuroprotective effects—potentially even surpassing those of IRF7—by limiting excessive inflammasome activation and restraining downstream neurotoxic signaling.

Targeted upregulation of ISG15 in microglia effectively inhibited microglial proliferation and induced a phenotypic shift toward an anti-inflammatory state. In line with previous findings, microglia exhibiting a highly ramified morphology are typically associated with homeostatic conditions, whereas amoeboid or rounded microglia with reduced branching complexity are indicative of an activated, proinflammatory state (Shui et al., 2024). Notably, overexpression of ISG15 resulted in a marked reduction in microglial density, an increased proportion of Arg1⁺ cells, and enhanced branching complexity, collectively suggesting suppression of proinflammatory activation. Mechanistically, this phenotype was accompanied by attenuated activation of the NLRP3 signaling pathway and improved neuronal survival within the ischemic hemisphere.

Stroke elicits a pronounced inflammatory response characterized by rapid microglial activation and directed migration toward the ischemic core. Increasing evidence indicates that modulating microglial motility can significantly influence post-stroke outcomes. For instance, PINK1 siRNA treatment has been reported to enhance microglial phagocytosis, migratory capacity, and anti-inflammatory polarization, with pretreatment effectively attenuating autophagy induction, reducing infarct volume, and ameliorating motor deficits in photothrombotic stroke models (Choi et al., 2023). Similarly, Matrix metalloproteinases have been shown to promote microglial and astrocytic migration, facilitating the formation of a protective barrier within the penumbral region and thereby limiting infarct expansion (X. Zhang, Zhao, Li, & Li, 2019). In contrast, other studies suggest that S100B augments M1 polarization and enhances microglial migration, ultimately exacerbating ischemic injury (S. Zhou, Zhu, Zhang, Pan, & Bao, 2018). Furthermore, Cannflavin A inhibits 2-arachidonoylglycerol-induced microglial migration has demonstrated therapeutic potential in experimental stroke paradigms (Jalin et al., 2015), whereas activation of Netrin-1 receptor signaling appears to attenuate microglial apoptosis and migration, contributing to neuroprotection in both human and rodent models (X. Yang, Liu, Zhong, Li, & Zhang, 2023). Collectively, these findings underscore the context-dependent and multifaceted roles of microglial migration in ischemic stroke and reperfusion injury. The present study revealed that siISG15-treated microglia displayed enhanced migratory activity following OGD/R exposure. Additional mechanistic investigations are needed to clarify how the increased migratory ability of microglial cells after knockdown of ISG15 contributes to ischemia–reperfusion.

In the present study, we observed that HGF/ISG15 attenuates NLRP3 inflammasome activation (Yu et al., 2023), although the precise mechanisms underlying this effect remain to be fully elucidated. ISG15, an IFN-induced ubiquitin-like modifier, covalently conjugates to lysine residues via a conserved C-terminal diglycine motif, a process known as ISG15ylation. This post-translational modification has been shown to regulate protein stability and interactions, often by antagonizing ubiquitination and proteasomal degradation (Kim & Yoo, 2010; Tecalco-Cruz & Zepeda-Cervantes, 2023). Consistent with previous reports, siRNA-mediated knockdown of ISG15 promoted NLRP3 degradation. Interestingly, however, NLRP3 expression was paradoxically increased in microglia subjected to OGD/R following siISG15 treatment. Further investigation revealed that ISG15 knockdown enhanced the mRNA stability of both NLRP3 and NF-кB, suggesting a dual regulatory role at the post-transcriptional as well as post-translational levels. mRNA stability is tightly regulated by multiple factors, including RNA-binding proteins (Liu et al., 2024), microRNAs (Peng et al., 2020), and epigenetic modifications such as N6-methyladenosine (m6A). However, the precise mechanisms by which ISG15 modulates mRNA turnover remain unclear. Our RNA-seq analysis suggests that this effect may involve genes implicated in RNA splicing and processing, raising the possibility that ISG15 influences mRNA fate through post-transcriptional regulatory pathways. Given that ISG15ylation exhibits selectivity for medium-sized proteins (Nakka et al., 2011), it is plausible that ISG15ylation may also display substrate specificity. Identification of the direct molecular targets of ISG15 in ischemic stroke will therefore be an important focus for future studies.

Despite these advances, several limitations of the current study should be acknowledged. While we demonstrated that ISG15 improves motor function and mitigates neurological deficits following ischemic stroke, its effects on cognitive, emotional, and memory-related outcomes were not evaluated. Moreover, ISG15 exists in both conjugated and free forms, and the relative contribution of each form to its neuroprotective effects remains unclear.

## 5. CONCLUSION

In conclusion, our findings indicate that both IRF7 and ISG15 exert neuroprotective effects in the context of ischemic stroke. Notably, microglial overexpression of ISG15 attenuates ischemia-induced brain injury by modulating inflammasome activation and stabilizing inflammation-related mRNAs. These results highlight ISG15 as a promising therapeutic target and provide a foundation for future mechanistic studies aimed at dissecting its role in post-stroke neuroinflammation.

## List of abbreviations

OGD/R: oxygen-glucose deprivation/reoxygenation
BBB: blood-brain barrier
NF-κB: nuclear factor-κB
TLRs: Toll-like receptors
IRFs: interferon regulatory factors
LPS: lipopolysaccharide
CpG-ODN: CpG oligodeoxynucleotides
tMCAO: transient middle cerebral artery occlusion
CCA: right common carotid artery
ECA: external carotid artery
ICA: internal carotid artery
MCA: middle cerebral artery
TTC: triphenyltetrazolium chloride
PFA: paraformaldehyde
PMSF: phenylmethylsulfonyl fluoride
TBST: Tris-buffered saline with Tween 20
NCM: normal culture medium
PS: Penicillin-Streptomycin
PPI: Protein-protein interaction
ISG15⁻/⁻: ISG15 knockout
HGF: hepatocyte growth factor
DMEM: Dulbecco’s Modified Eagle’s Medium
FBS: fetal bovine serum
DEGs: Differentially expressed genes
GO: Gene Ontology
KEGG: Kyoto Encyclopedia of Genes and Genomes
GSEA: Gene set enrichment analysis
SD: standard deviation
pIRF7: phosphorylated
IRF7 Il-1βmat: active form of Il-1β
GSDMDmat: active form of GSDMD
GSDMDpre: precursor of GSDMD
NC: negative control
CHX: cycloheximide
TBI: traumatic brain injury
siISG15: siRNA targeting ISG15
siIRF7: siRNA targeting IRF7

## ACKNOWLEDGMENTS

This research was funded by the Sichuan Provincial Science and Technology Program Project (2023NSFSC0643, 2023YFS0050, 2024YFHZ0009).

## Reference

1. Choi, S. G., Shin, J., Lee, K. Y., Park, H., Kim, S. I., Yi, Y. Y., . . . Shin, H. J. (2023). PINK1 siRNA-loaded poly(lactic-co-glycolic acid) nanoparticles provide neuroprotection in a mouse model of photothrombosis-induced ischemic stroke. Glia, 71(5), 1294–1310. doi:10.1002/glia.24339

2. Chong, M., Sjaarda, J., Pigeyre, M., Mohammadi-Shemirani, P., Lali, R., Shoamanesh, A., . . . Pare, G. (2019). Novel Drug Targets for Ischemic Stroke Identified Through Mendelian Randomization Analysis of the Blood Proteome. Circulation, 140(10), 819–830. doi:10.1161/CIRCULATIONAHA.119.040180

3. Clarkson, B. D. S., Grund, E., David, K., Johnson, R. K., & Howe, C. L. (2022). ISGylation is induced in neurons by demyelination driving ISG15-dependent microglial activation. J Neuroinflammation, 19(1), 258. doi:10.1186/s12974-022-02618-4

4. Cohen, M., Matcovitch, O., David, E., Barnett-Itzhaki, Z., Keren-Shaul, H., Blecher-Gonen, R., . . . Schwartz, M. (2014). Chronic exposure to TGFbeta1 regulates myeloid cell inflammatory response in an IRF7-dependent manner. EMBO J, 33(24), 2906–2921. doi:10.15252/embj.201489293

5. Das, A., Arifuzzaman, S., Yoon, T., Kim, S. H., Chai, J. C., Lee, Y. S., . . . Chai, Y. G. (2017). RNA sequencing reveals resistance of TLR4 ligand-activated microglial cells to inflammation mediated by the selective jumonji H3K27 demethylase inhibitor. Sci Rep, 7(1), 6554. doi:10.1038/s41598-017-06914-5

6. De Simoni, M. G., Storini, C., Barba, M., Catapano, L., Arabia, A. M., Rossi, E., & Bergamaschini, L. (2003). Neuroprotection by complement (C1) inhibitor in mouse transient brain ischemia. J Cereb Blood Flow Metab, 23(2), 232–239. doi:10.1097/01.WCB.0000046146.31247.A1

7. Deng, X., Hu, Z., Zhou, S., Wu, Y., Fu, M., Zhou, C., . . . Huang, Y. (2024). Perspective from single-cell sequencing: Is inflammation in acute ischemic stroke beneficial or detrimental? CNS Neurosci Ther, 30(1), e14510. doi:10.1111/cns.14510

8. Dibas, J., Al-Saad, H., & Dibas, A. (2019). Basics on the use of acid-sensing ion channels’ inhibitors as therapeutics. Neural Regen Res, 14(3), 395–398. doi:10.4103/1673-5374.245466

9. Gao, D., Hao, J. P., Li, B. Y., Zheng, C. C., Miao, B. B., Zhang, L., . . . Zhang, L. (2023). Tetrahydroxy stilbene glycoside ameliorates neuroinflammation for Alzheimer’s disease via cGAS-STING. Eur J Pharmacol, 953, 175809. doi:10.1016/j.ejphar.2023.175809

10. Gao, Y., Lu, X., Zhang, G., Liu, C., Sun, S., Mao, W., . . . Zhang, L. (2024). DRD4 alleviates acute kidney injury by suppressing ISG15/NOX4 axis-associated oxidative stress. Redox Biol, 70, 103078. doi:10.1016/j.redox.2024.103078

11. Hain, H. S., Pandey, R., Bakay, M., Strenkowski, B. P., Harrington, D., Romer, M., . . . Hakonarson, H. (2021). Inducible knockout of Clec16a in mice results in sensory neurodegeneration. Sci Rep, 11(1), 9319. doi:10.1038/s41598-021-88895-0

12. He, C., Li, Y., Chen, Z. Y., & Huang, C. K. (2023). Crosstalk of renal cell carcinoma cells and tumor-associated macrophages aggravates tumor progression by modulating muscleblind-like protein 2/B-cell lymphoma 2/beclin 1-mediated autophagy. Cytotherapy, 25(3), 298–309. doi:10.1016/j.jcyt.2022.09.001

13. Ho, M. H., Yen, C. H., Hsieh, T. H., Kao, T. J., Chiu, J. Y., Chiang, Y. H., . . . Chou, S. Y. (2021). CCL5 via GPX1 activation protects hippocampal memory function after mild traumatic brain injury. Redox Biol, 46, 102067. doi:10.1016/j.redox.2021.102067

14. Hu, Y., Hong, X. Y., Yang, X. F., Ma, R. H., Wang, X., Zhang, J. F., . . . Liu, G. P. (2019). Inflammation-dependent ISG15 upregulation mediates MIA-induced dendrite damages and depression by disrupting NEDD4/Rap2A signaling. Biochim Biophys Acta Mol Basis Dis, 1865(6), 1477–1489. doi:10.1016/j.bbadis.2019.02.020

15. Huang, J., Lee, H. Y., Zhao, X., Han, J., Su, Y., Sun, Q., . . . Dong, C. (2021). Interleukin -17D regulates group 3 innate lymphoid cell function through its receptor CD93. Immunity, 54(4), 673–686 e674. doi:10.1016/j.immuni.2021.03.018

16. Huang, Y., Happonen, K. E., Burrola, P. G., O’Connor, C., Hah, N., Huang, L., . . . Lemke, G. (2021). Microglia use TAM receptors to detect and engulf amyloid beta plaques. Nat Immunol, 22(5), 586–594. doi:10.1038/s41590-021-00913-5

17. Jalin, A. M., Rajasekaran, M., Prather, P. L., Kwon, J. S., Gajulapati, V., Choi, Y., . . . Kim, W. K. (2015). Non-Selective Cannabinoid Receptor Antagonists, Hinokiresinols Reduce Infiltration of Microglia/Macrophages into Ischemic Brain Lesions in Rat via Modulating 2 - Arachidonolyglycerol-Induced Migration and Mitochondrial Activity. PLoS One, 10(10), e0141600. doi:10.1371/journal.pone.0141600

18. Jeon, E., Seo, M. S., Lkhagva-Yondon, E., Lim, Y. R., Kim, S. W., Kang, Y. J., . . . Jeon, M. S. (2024). Neuroprotective effect of L-DOPA-induced interleukin-13 on striatonigral degeneration in cerebral ischemia. Cell Death Dis, 15(11), 854. doi:10.1038/s41419-024-07252-x

19. Kang, J. A., Kim, Y. J., & Jeon, Y. J. (2022). The diverse repertoire of ISG15: more intricate than initially thought. Exp Mol Med, 54(11), 1779–1792. doi:10.1038/s12276-022-00872-3

20. Khorooshi, R., & Owens, T. (2010). Injury-induced type I IFN signaling regulates inflammatory responses in the central nervous system. J Immunol, 185(2), 1258–1264. doi:10.4049/jimmunol.0901753

21. Kim, M. J., & Yoo, J. Y. (2010). Inhibition of hepatitis C virus replication by IFN-mediated ISGylation of HCV-NS5A. J Immunol, 185(7), 4311–4318. doi:10.4049/jimmunol.1000098

22. Kumar, R., Palmer, E., Gardner, A. E., Carroll, R., Banka, S., Abdelhadi, O., . . . Gecz, J. (2020). Expanding Clinical Presentations Due to Variations in THOC2 mRNA Nuclear Export Factor. Front Mol Neurosci, 13, 12. doi:10.3389/fnmol.2020.00012

23. Lenart, N., Cserep, C., Csaszar, E., Posfai, B., & Denes, A. (2024). Microglia-neuron-vascular interactions in ischemia. Glia, 72(5), 833–856. doi:10.1002/glia.24487

24. Li, L., Ulrich, R., Baumgartner, W., & Gerhauser, I. (2015). Interferon -stimulated genes-essential antiviral effectors implicated in resistance to Theiler’s virus-induced demyelinating disease. J Neuroinflammation, 12, 242. doi:10.1186/s12974-015-0462-x

25. Li, L., Yang, J. H., Li, C., Zhou, H. F., Yu, L., Wu, X. L., . . . Wan, H. T. (2023). Danhong injection improves neurological function in rats with ischemic stroke by enhancing neurogenesis and activating BDNF/AKT/CREB signaling pathway. Biomed Pharmacother, 163, 114887. doi:10.1016/j.biopha.2023.114887

26. Li, Z., Huang, Q., Chen, H., Lin, Z., Zhao, M., & Jiang, Z. (2017). Interferon Regulatory Factor 7 Promoted Glioblastoma Progression and Stemness by Modulating IL-6 Expression in Microglia. J Cancer, 8(2), 207–219. doi:10.7150/jca.16415

27. Liang, Y. B., Guo, Y. Q., Song, P. P., Zhu, Y. H., Zhu, P. Z., Liu, R. R., . . . Zhang, Y. S. (2020). Memantine ameliorates tau protein deposition and secondary damage in the ipsilateral thalamus and sensory decline following focal cortical infarction in rats. Neurosci Lett, 731, 135091. doi:10.1016/j.neulet.2020.135091

28. Liang, Z., Lou, Y., Hao, Y., Li, H., Feng, J., & Liu, S. (2023). The Relationship of Astrocytes and Microglia with Different Stages of Ischemic Stroke. Curr Neuropharmacol, 21(12), 2465–2480. doi:10.2174/1570159X21666230718104634

29. Liu, C., Chen, H., Tao, X., Li, C., Li, A., & Wu, W. (2024). ALKBH5 protects against stroke by reducing endoplasmic reticulum stress-dependent inflammation injury via the STAT5/PERK/EIF2alpha/CHOP signaling pathway in an m(6)A-YTHDF1-dependent manner. Exp Neurol, 372, 114629. doi:10.1016/j.expneurol.2023.114629

30. Nakka, V. P., Lang, B. T., Lenschow, D. J., Zhang, D. E., Dempsey, R. J., & Vemuganti, R. (2011). Increased cerebral protein ISGylation after focal ischemia is neuroprotective. J Cereb Blood Flow Metab, 31(12), 2375–2384. doi:10.1038/jcbfm.2011.103

31. Orsini, F., Villa, P., Parrella, S., Zangari, R., Zanier, E. R., Gesuete, R., . . . De Simoni, M. G. (2012). Targeting mannose-binding lectin confers long-lasting protection with a surprisingly wide therapeutic window in cerebral ischemia. Circulation, 126(12), 1484–1494. doi:10.1161/CIRCULATIONAHA.112.103051

32. Packer, J. M., Bray, C. E., Beckman, N. B., Wangler, L. M., Davis, A. C., Goodman, E. J., . . . Godbout, J. P. (2024). Impaired cortical neuronal homeostasis and cognition after diffuse traumatic brain injury are dependent on microglia and type I interferon responses. Glia, 72(2), 300–321. doi:10.1002/glia.24475

33. Pappas, D. J., Gabatto, P. A., Oksenberg, D., Khankhanian, P., Baranzini, S. E., Gan, L., & Oksenberg, J. R. (2012). Transcriptional expression patterns triggered by chemically distinct neuroprotective molecules. Neuroscience, 226, 10–20. doi:10.1016/j.neuroscience.2012.09.007

34. Patil, S., Rossi, R., Jabrah, D., & Doyle, K. (2022). Detection, Diagnosis and Treatment of Acute Ischemic Stroke: Current and Future Perspectives. Front Med Technol, 4, 748949. doi:10.3389/fmedt.2022.748949

35. Peng, Z., Li, M., Tan, X., Xiang, P., Wang, H., Luo, Y., . . . Yang, J. (2020). miR-211-5p alleviates focal cerebral ischemia-reperfusion injury in rats by down-regulating the expression of COX2. Biochem Pharmacol, 177, 113983. doi:10.1016/j.bcp.2020.113983

36. Penzes, M., Turos, D., Mathe, D., Szigeti, K., Hegedus, N., Rauscher, A. A., Malnasi-Csizmadia, A. (2020). Direct myosin-2 inhibition enhances cerebral perfusion resulting in functional improvement after ischemic stroke. Theranostics, 10(12), 5341–5356. doi:10.7150/thno.42077

37. Peters, V. A., Joesting, J. J., & Freund, G. G. (2013). IL-1 receptor 2 (IL-1R2) and its role in immune regulation. Brain Behav Immun, 32, 1–8. doi:10.1016/j.bbi.2012.11.006

38. Przanowski, P., Loska, S., Cysewski, D., Dabrowski, M., & Kaminska, B. (2018). ISG’ylation increases stability of numerous proteins including Stat1, which prevents premature termination of immune response in LPS-stimulated microglia. Neurochem Int, 112, 227–233. doi:10.1016/j.neuint.2017.07.013

39. Ranjbar, K., Komaki, A., Fayazi, B., & Zarrinkalam, E. (2024). Coenzyme Q10 and exercise training reinstate middle cerebral artery occlusion-induced behavioral deficits and hippocampal long-term potentiation suppression in aging rats. Psychopharmacology (Berl), 241(8), 1577–1594. doi:10.1007/s00213-024-06583-z

40. Salaudeen, M. A., Bello, N., Danraka, R. N., & Ammani, M. L. (2024). Understanding the Pathophysiology of Ischemic Stroke: The Basis of Current Therapies and Opportunity for New Ones. Biomolecules, 14(3). doi:10.3390/biom14030305

41. Shui, X., Chen, J., Fu, Z., Zhu, H., Tao, H., & Li, Z. (2024). Microglia in Ischemic Stroke: Pathogenesis Insights and Therapeutic Challenges. J Inflamm Res, 17, 3335–3352. doi:10.2147/JIR.S461795

42. Somebang, K., Rudolph, J., Imhof, I., Li, L., Niemi, E. C., Shigenaga, J., . . . Hsieh, C. L. (2021). CCR2 deficiency alters activation of microglia subsets in traumatic brain injury. Cell Rep, 36(12), 109727. doi:10.1016/j.celrep.2021.109727

43. Stevens, S. L., Leung, P. Y., Vartanian, K. B., Gopalan, B., Yang, T., Simon, R. P., & Stenzel-Poore, M. P. (2011). Multiple preconditioning paradigms converge on interferon regulatory factor-dependent signaling to promote tolerance to ischemic brain injury. J Neurosci, 31(23), 8456–8463. doi:10.1523/JNEUROSCI.0821-11.2011

44. Tanaka, T., Murakami, K., Bando, Y., & Yoshida, S. (2015). Interferon regulatory factor 7 participates in the M1-like microglial polarization switch. Glia, 63(4), 595–610. doi:10.1002/glia.22770

45. Tecalco-Cruz, A. C., & Zepeda-Cervantes, J. (2023). Protein ISGylation: a posttranslational modification with implications for malignant neoplasms. Explor Target Antitumor Ther, 4(4), 699–715. doi:10.37349/etat.2023.00162

46. van Olst, L., Simonton, B., Edwards, A. J., Forsyth, A. V., Boles, J., Jamshidi, P., . . . Gate, D. (2025). Microglial mechanisms drive amyloid-beta clearance in immunized patients with Alzheimer’s disease. Nat Med. doi:10.1038/s41591-025-03574-1

47. Wang, T., Liu, Q., & Duan, L. (2020). MBNL1 regulates resistance of HeLa cells to cisplatin via Nrf2. Biochem Biophys Res Commun, 522(3), 763–769. doi:10.1016/j.bbrc.2019.11.162

48. Weichert, L., Dusedau, H. P., Fritzsch, D., Schreier, S., Scharf, A., Grashoff, M., . . . Kroger, A. (2023). Astrocytes evoke a robust IRF7-independent type I interferon response upon neurotropic viral infection. J Neuroinflammation, 20(1), 213. doi:10.1186/s12974-023-02892-w

49. Wu, H., Peng, B., Mohammed, F. S., Gao, X., Qin, Z., Sheth, K. N., . . . Jiang, Z. (2022). Brain Targeting, Antioxidant Polymeric Nanoparticles for Stroke Drug Delivery and Therapy. Small, 18(22), e2107126. doi:10.1002/smll.202107126

50. Xie, C., Long, F., Li, L., Li, X., Ma, M., Lu, Z., . . . Lin, C. (2022). PTBP3 modulates P53 expression and promotes colorectal cancer cell proliferation by maintaining UBE4A mRNA stability. Cell Death Dis, 13(2), 128. doi:10.1038/s41419-022-04564-8

51. Yang, H., Lin, C. H., Ma, G., Baffi, M. O., & Wathelet, M. G. (2003). Interferon regulatory factor-7 synergizes with other transcription factors through multiple interactions with p300/CBP coactivators. J Biol Chem, 278(18), 15495–15504. doi:10.1074/jbc.M212940200

52. Yang, X., Liu, Y., Zhong, W., Li, Y., & Zhang, W. (2023). Netrin-1 controls inflammation in response to ischemic stroke through altering microglia phenotype. Front Immunol, 14, 1178638. doi:10.3389/fimmu.2023.1178638

53. Yang, Y., Zhang, M., Li, N., Wang, C., Yang, H., Hou, X., . . . Wu, K. (2024). Hirschsprung’s disease: m6A methylase VIRMA suppresses cell migration and proliferation by regulating GSK3beta. Pediatr Res, 96(4), 942–951. doi:10.1038/s41390-024-03136-0

54. Yu, L., Zhang, Z., Chen, H., Wang, M., Mao, W., Hu, J., . . . Cui, G. (2023). Remote limb ischemic postconditioning inhibits microglia pyroptosis by modulating HGF after acute ischemia stroke. Bioeng Transl Med, 8(6), e10590. doi:10.1002/btm2.10590

55. Zhang, K., Zhang, Y., Li, Z., Chen, J., Chang, Y., Li, Y., . . . Huang, K. (2025). Potentiating microglial efferocytosis by MFG-E8 improves survival and neurological outcome after successful cardiopulmonary resuscitation in mice. Brain Pathol, 35(4), e13327. doi:10.1111/bpa.13327

56. Zhang, X., Zhao, H. H., Li, D., & Li, H. P. (2019). Neuroprotective effects of matrix metalloproteinases in cerebral ischemic rats by promoting activation and migration of astrocytes and microglia. Brain Res Bull, 146, 136–142. doi:10.1016/j.brainresbull.2018.11.003

57. Zhao, P., Hou, L., Farley, K., Sundrud, M. S., & Remold-O’Donnell, E. (2014). SerpinB1 regulates homeostatic expansion of IL-17+ gammadelta and CD4+ Th17 cells. J Leukoc Biol, 95(3), 521–530. doi:10.1189/jlb.0613331

58. Zhao, Y., Li, Q., Niu, J., Guo, E., Zhao, C., Zhang, J., . . . Yang, K. (2024). Neutrophil Membrane-Camouflaged Polyprodrug Nanomedicine for Inflammation Suppression in Ischemic Stroke Therapy. Adv Mater, 36(21), e2311803. doi:10.1002/adma.202311803

59. Zhou, L., Jin, Y., Wu, D., Cun, Y., Zhang, C., Peng, Y., Zhang, P. (2023). Current evidence, clinical applications, and future directions of transcranial magnetic stimulation as a treatment for ischemic stroke. Front Neurosci, 17, 1177283. doi:10.3389/fnins.2023.1177283

60. Zhou, S., Zhu, W., Zhang, Y., Pan, S., & Bao, J. (2018). S100B promotes microglia M1 polarization and migration to aggravate cerebral ischemia. Inflamm Res, 67(11-12), 937–949. doi:10.1007/s00011-018-1187-y

61. Zhou, Y., Pan, J., Peng, Q., Dong, Z., Deng, L., & Wang, Y. (2017). The TLR9 Antagonist iCpG-ODN at Different Dosages Inhibits Cerebral Ischemia/Reperfusion Injury in Mice. CNS Neurol Disord Drug Targets, 16(5), 624–633. doi:10.2174/1871527316666170206150259

