## Supplementary Tables and Figures for "Activation of IRF7/ISG15 axis in microglia inhibits NLRP3 Expression and Improves the Prognosis of Ischemia/Reperfusion in Mice"

**Supplementary Table 1. Modified DeSimoni neuroscore for the assessment of general deficits (De Simoni et al., 2003; Orsini et al., 2012)**

| **Objective** | **Assessment/instruction** | **Score=0** | **Score=1** | **Score=2** | **Score=3** | **Score=4** |
| --- | --- | --- | --- | --- | --- | --- |
| Hair | Mouse observed on open bench top (OS) without interference | Hair neat and clean | Lack of grooming, piloerection and dirt on the fur around nose and eyes | Lack of grooming, piloerection, and dirty coat, extending beyond just nose and eyes |  |  |
| Eyes | Mouse on OS without interference or stimulation | Open and clear | Open and characterized by milky white mucus | Open and characterized by milky dark mucus | Eyes clotted (one or both  sides) | Closed |
| Posture | Place the mouse on the palm of your hand and rock gently to observe stability | The mouse stands in the upright position on four limbs with the back parallel to the palm. During the rocking movement, it uses its limbs to stabilize itself | The mouse stands humpbacked. During the rocking movement, it lowers its body instead of using its limbs to gain stability | The head or part of the trunk lies on the palm | The mouse reclines to one side but may be able to turn to an upright position with some difficulty | No upright position possible |
| Spontaneous Activity | Mouse on OS without interference or stimulation | The mouse is alert and explores actively | The mouse seems alert, but it is calm and quiet | The mouse starts and stops exploring repeatedly and slowly. The mouse is listless, moves sluggishly but does not explore | The mouse is lethargic or stuporous and barely moves during the 60s | No spontaneous movements |

**Supplementary Table 2. Modified DeSimoni neuroscore for the assessment of focal deficits (De Simoni et al., 2003; Orsini et al., 2012)**

| **Objective** | **Assessment/instruction** | **Score=0** | **Score=1** | **Score=2** | **Score=3** | **Score=4** |
| --- | --- | --- | --- | --- | --- | --- |
| Body symmetry | Mouse on OS, observation of undisturbed resting behaviour and description of the virtual nose–tail line | Normal. a, Body: normal posture,trunk elevated from the bench,with forelimbs and hindlimbs leaning beneath the body. b,Tail: straight | Slight asymmetry. a, Body: leans on one side with forelimbs and hindlimbs  leaning beneath the body. b, Tail: slightly bent | Moderate asymmetry. a, Body: leans on one side with forelimbs and hindlimbs stretched out. b, Tail: slightly bent | Clear asymmetry. a, Body leans on one. b, Tail: clearly bent | Complete asymmetry. a, Body. b, Tail |
| Gait | Mouse on OS. Observation of undisturbed movements | Normal. Gait is flexible, symmetric,and quick | Stiff, inflexible. The mouse walks humpbacked,slower than normal mice | Limping with asymmetric movements | More severe limping, drifting,falling with obvious deficiency in gait | Does not walk spontaneously.  (In this case, stimulation  will be performed gently  pushing the mouse with  a pen. When stimulated,  the mouse walks no  longer than three steps.) |
| Climbing | Mouse is placed in the center of a  gripping surface at an angle of  45° to OS | Normal. The mouse climbs quickly | Climbs slowly, limb weakness  present | Holds onto slope, does not slip  or climb | Slides down slope; difficulty to  prevent fall | Slides down slope, unsuccessful  effort to prevent fall |
| Circling behavior | Mouse on OS. Observation of the  mouse walking undisturbed on  the OS | Circling behavior absent. The  mouse turns equally to left or  right | Predominantly one-sided  turns. Optional: record to  which side the mouse  turns | Circles to one side, although  not constantly | Circles constantly to one side.  This one is now highlighted  in yellow | No movements |
| Forelimb symmetry | Mouse suspended by the tail.  Movements and position of  forelimbs are observed | Normal. Both forelimbs are extended  towards the bench and  move actively | Light asymmetry. Contralateral  forelimb does not  extend entirely | Marked asymmetry. Contralateral  forelimb bends towards  the trunk. The body  slightly bends on the side  ipsilateral to the stroke | Prominent asymmetry. Contralateral  forelimb adheres  to the trunk | Slight asymmetry, no body/  limb movement |
| Compulsory circling | Forelimbs on bench, hindlimbs  suspended by the tail. This  position reveals the presence of  the contralateral limb palsy. In  this handstand position, limb  weakness is displayed by a  circling behavior when the  animal attempts forward motion | Absent. Normal extension of both  forelimbs | Both forelimbs extended but  begins to circle predominantly  to one side | Circles only to one side and  may fall to one side | Pivots to one side sluggishly  and does not rotate in a  full circle. Mouse will fall  to one side | No or rare movements |
| Gripping of the forepaw | Mouse is held by the tail on the  wire bar cage lid, so that the  forepaws touch the grid | Mouse grasps the grid firmly with  forepaws and tries to place the  hind paws also onto the grid  by pulling the hindpaws under  the body | Mouse accesses the grid but  has less power. A slight  pull breaks the grip of the  forepaws | Mouse cannot grip with the  impaired forepaw | Mouse cannot grip the grid |  |

**Supplementary Table 3. Antibodies used in this study.**

| **Antibody** | **WB/IF/Dilution** | **Sources** | **Catalog No** |
| --- | --- | --- | --- |
| mouse anti-IRF7 | IF，1:200 | Santa Cruz | sc-74471 |
| mouse anti-ISG15 | IF，1:200 | Santa Cruz | sc-166755 |
| Rabbit anti-IBA1  mouse anti-NeuN  Rabbit anti-GFAP  Mouse anti-Arg1  Goat Anti-Rabbit IgG H&L  Goat Anti-Mouse IgG H&L  Donkey Anti-Rabbit IgG H&L  Goat Anti-Mouse IgG H&L  Rabbit anti-phospho-IRF7  mouse anti-ISG15  mouse anti-NLRP3  Rabbit anti-NF-κB p65  Mouse anti-GSDMD  Mouse anti-IL-1β | IF，1:1000  IF，1:1000  IF，1:5000  IF，1:1000  IF，1:1000  IF，1:1000  IF，1:1000  IF，1:1000  WB，1:1000  WB，1:500  WB，1:1000  WB，1:10000  WB，1:100  WB，1:1000 | WAKO  Abcam  Abcam  Proteintech  Abcam  Abcam  Abcam  Abcam  BIOSS  Santa Cruz  Adipogen  Proteintech  Santa Cruz  Cell Signaling | 019-19741  ab279295  ab7260  66129-1-Ig  ab150077  ab150113  ab150075  ab150115  bs-3196R  sc-166755  AG-20B-0014-C100  80979-1-RR  sc-393581  12242 |

**Supplementary Table 4. The sequences of primers and siRNAs used in this study.**

| **Gene name** | **Forward primer 5’-3’** | **Reverse primer 5’-3’** |
| --- | --- | --- |
| GAPDH | CCCATCACCATCTTCCAGGAG | TTCACCACCTTCTTCTTGATGTCAT |
| NLRP3 | ATTACCCGCCCGAGAAAGG | TCGCAGCAAAGATCCACACAG |
| NF-κB | TCCTGTTCGAGTCTCCATGCAG | GGTCTCATAGGTCCTTTTGCGC |
| IL-1β | CAGGCTCCGAGATGAAC | TGCTTGTGAGGTGCTGA |
| ISG15  IRF7  GSDMD  Arg1  TNF-α  TGF-β  siIRF7  siISG15 | GGTGTCCGTGACTAACTCCAT  GCTATTGGGGGAGGTCAGCA  CCATCGGCCTTTGAGAAAGTG  GCTCAGGTGAATCGGCCTTT  CCCTCACACTCAGATCATCTTCT  CTCCCGTGGCTTCTAGTGC  CTTCGACTTCAGCACTTTCTT  GCACAGTGATGCTAGTGGT | TGGAAAGGGTAAGACCGTCCT  AACGCCCTGTGCTGTGGAG  ACACATGAATAACGGGGTTTCC  TGGCTTGCGAGACGTAGAC  GCTACGACGTGGGCTACAG  GCCTTAGTTTGGACAGGATCTG |

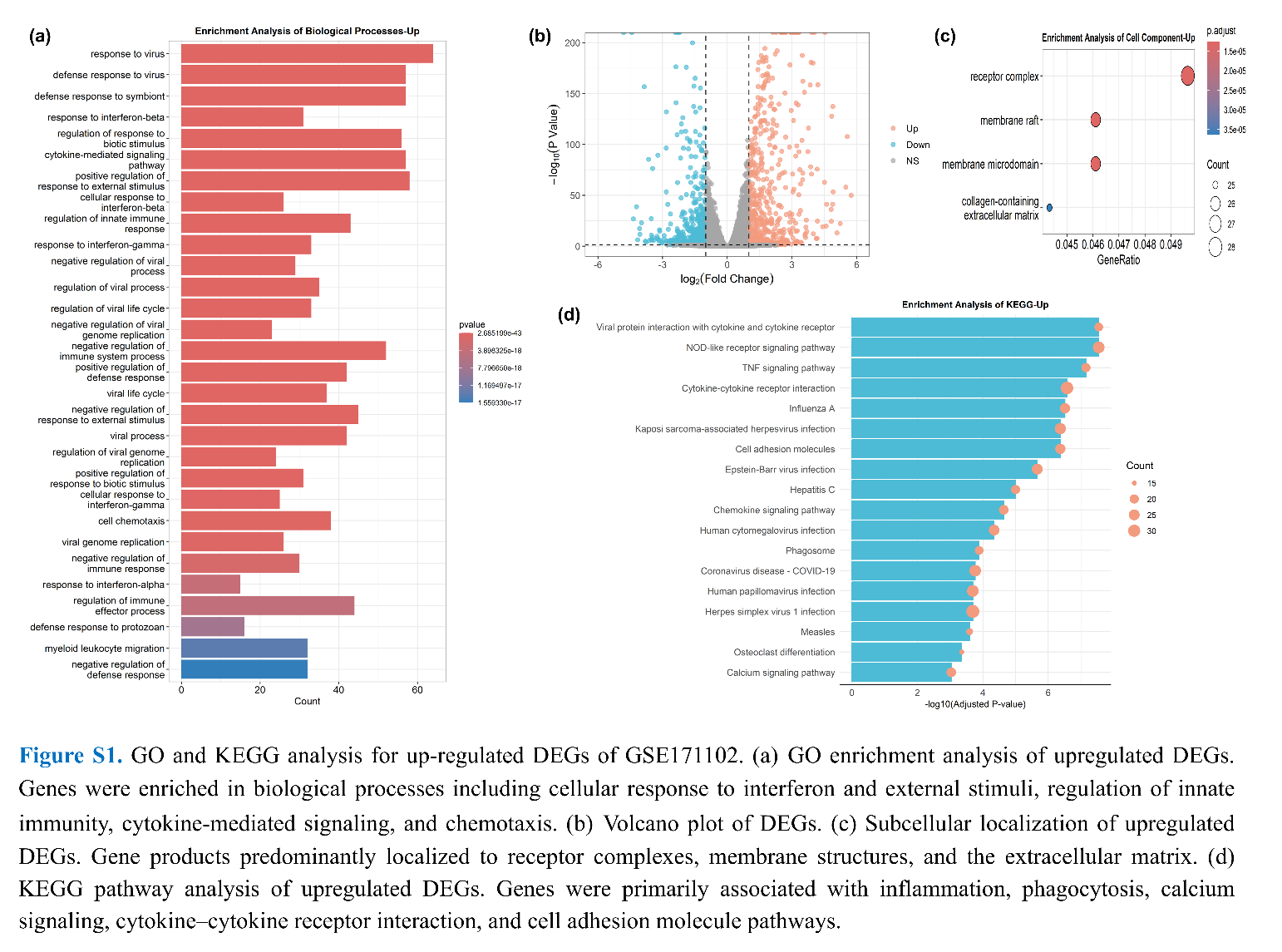

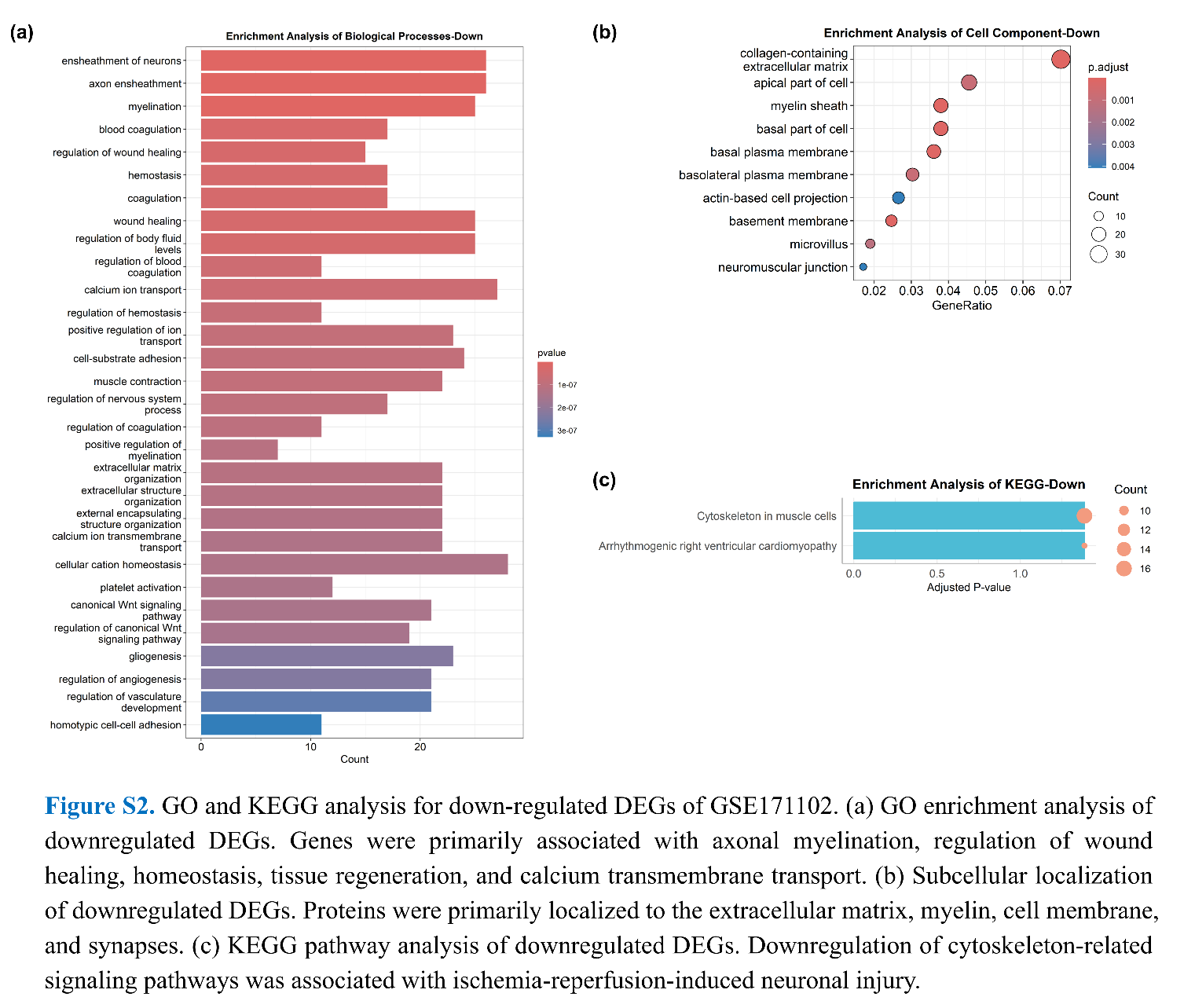

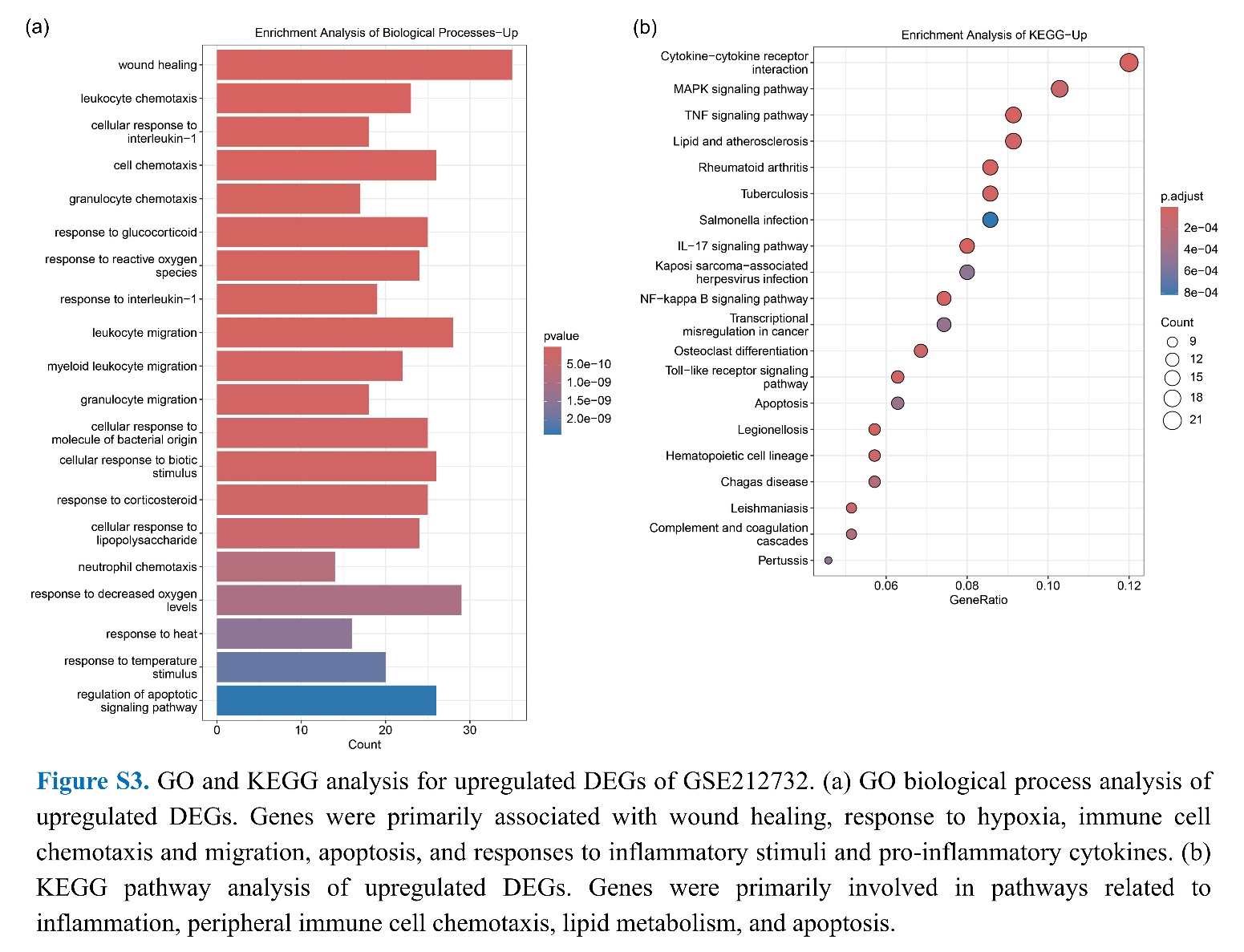

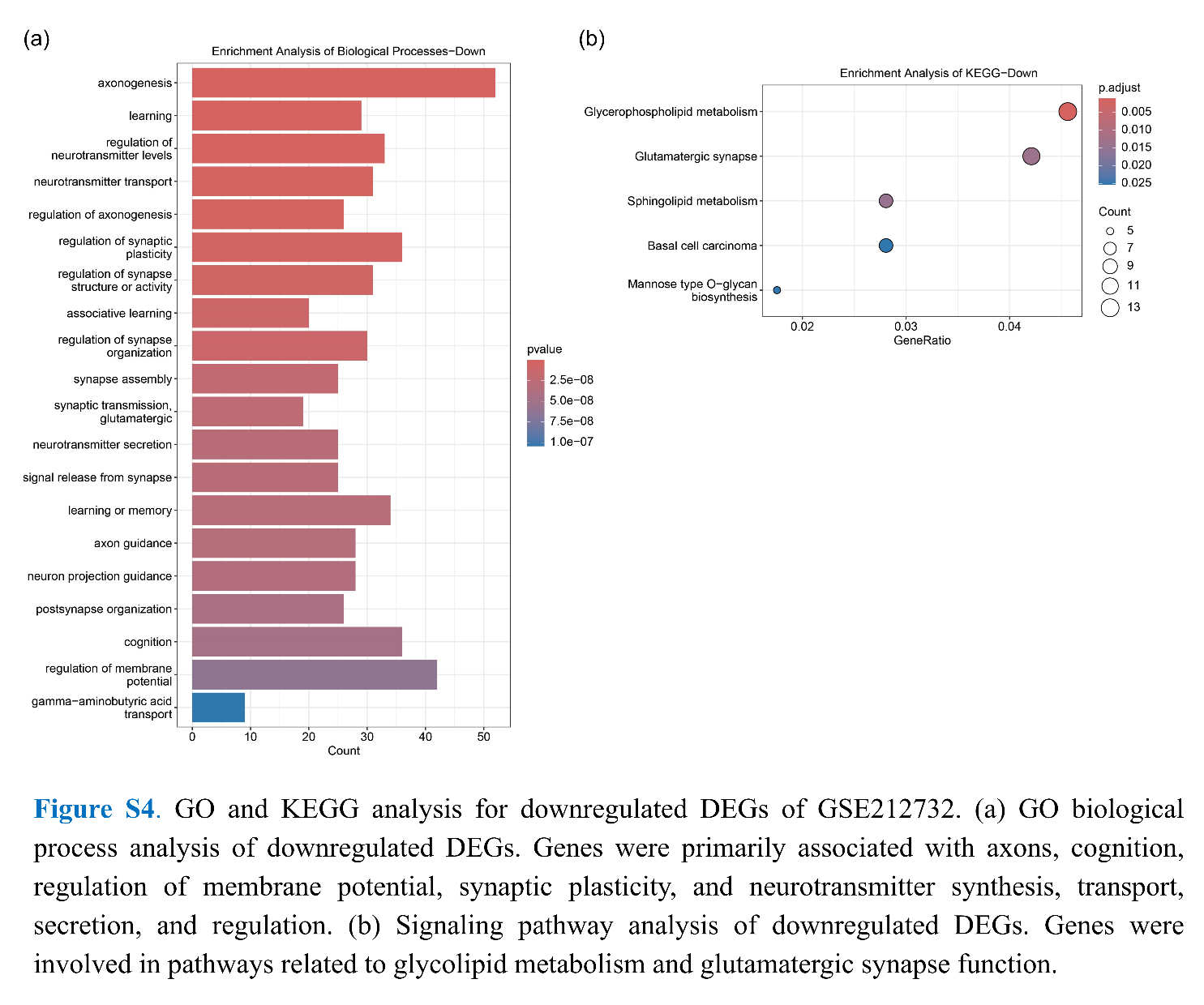

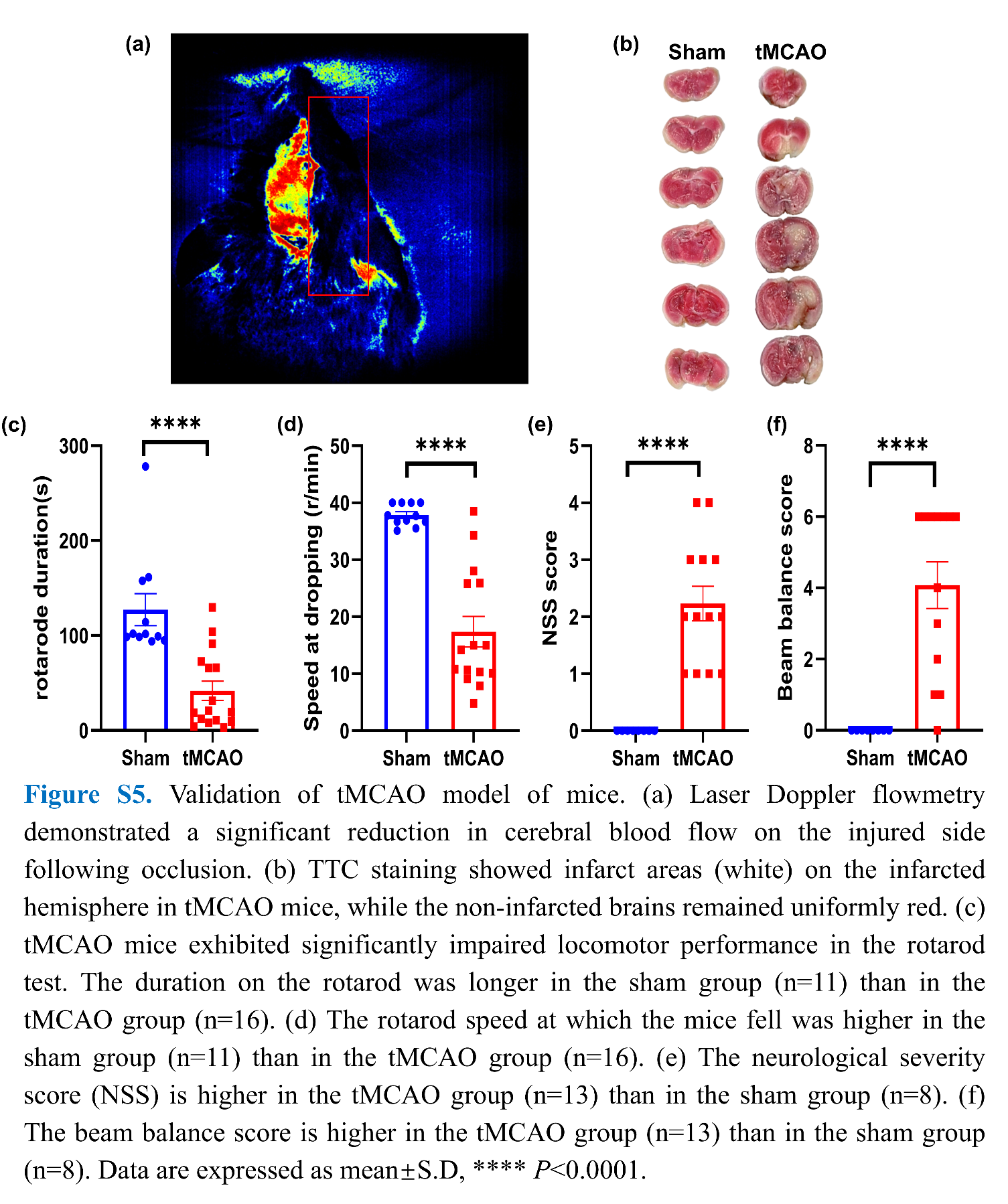

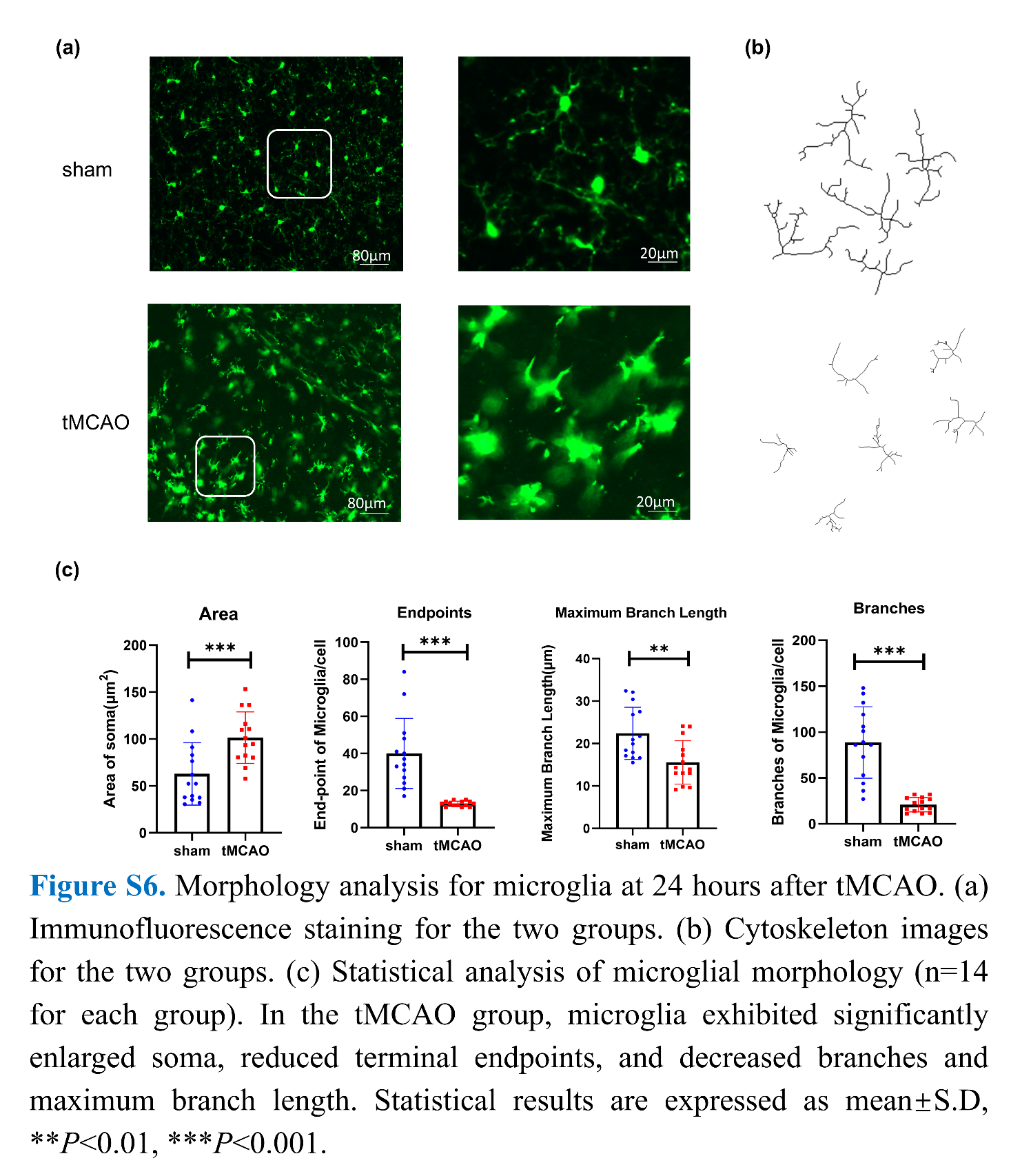

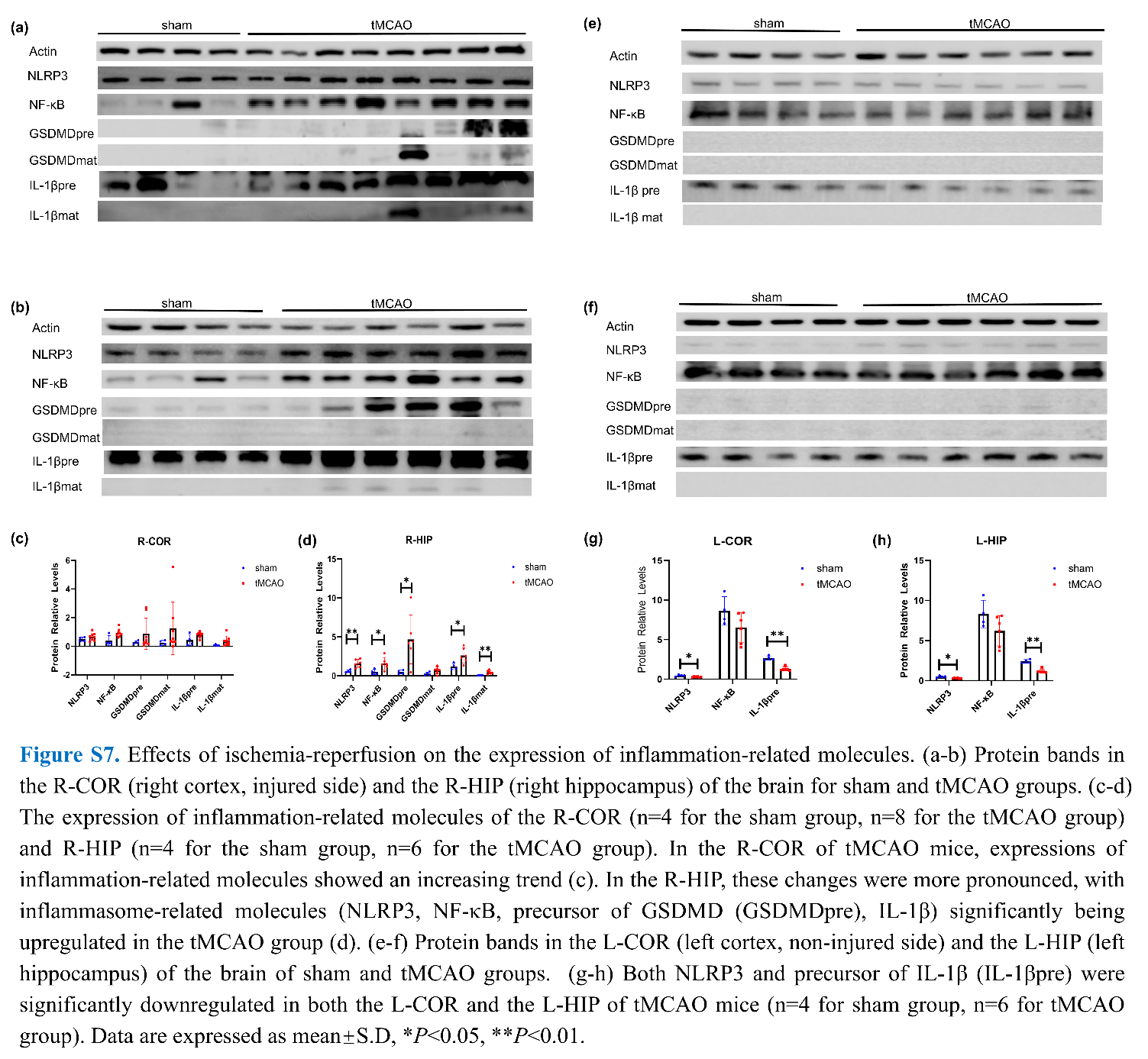

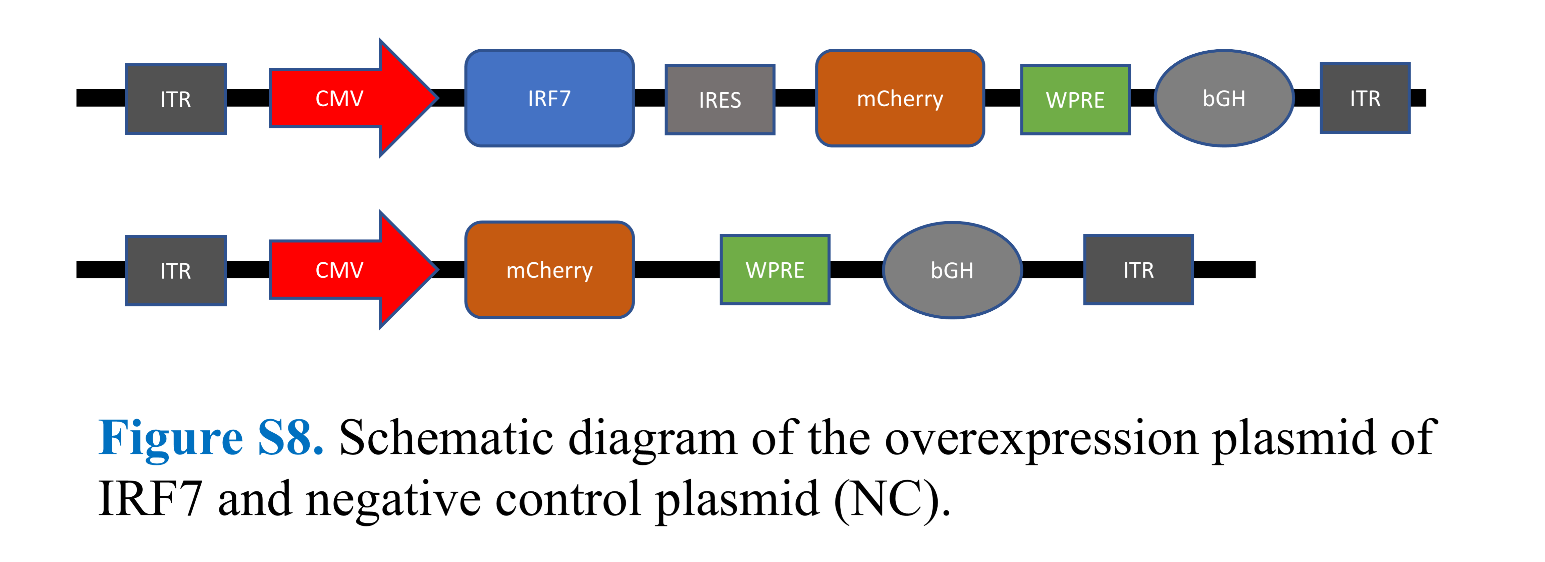
